# Counterintuitive Binding of Phosphorylated DEP Domain from Dishevelled Protein to Negatively Charged Membranes

**DOI:** 10.1101/2023.01.27.525887

**Authors:** Francesco L. Falginella, Martina Drabinová, Vítezslav Bryja, Robert Vácha

## Abstract

To accomplish its role of signaling hub in all Wnt signaling pathways, Dishevelled (DVL) protein needs to dynamically relocalize to the inner leaflet of the cellular plasma membrane (PM). Combined experimental and computational evidence showed that the binding of DVL to the PM is mainly driven by the electrostatic attraction between a stretch of positively charged amino acids located on the C-terminal DEP domain of DVL and anionic phospholipid species, with a striking preference for phosphatidic acid (PA). Here, by means of computational simulations and QCM-D experiments, we demonstrate that four recently identified phosphorylation sites on DEP domain, alter the electrostatic potential of the membrane binding interface, but do not prevent the recruitment to anionic membranes. On the contrary, the phosphorylated residues are involved in hydrogen bond and ion-mediated interactions with the lipid headgroup of PA. Our results suggest that the effect of phosphorylation on protein-membrane association could be counterintuitive and sensitive to changes in the local environment including specific lipids, salts, and pH.

**SIGNIFICANCE:** Phosphorylation regulates the cellular activity and localization of many peripheral proteins by, among others, decreasing the affinity for negatively charged membranes. Here, we report how phosphorylation affects the membrane interaction of DEP domain from Dishevelled protein, the intracellular signaling hub in Wnt pathways. We found that despite the negative charge induced by phosphorylation, DEP domain was steadily adsorbed to the surface of negatively charged PA-rich membranes, due to a dense network of cation-mediated interactions and hydrogen bonds.

## INTRODUCTION

Protein-membrane interactions represent a key step in the activation and progression of many cell signaling pathways (1, 2), among which Wnt signaling plays an indispensable role as it regulates cellular growth and differentiation during both tissue development and homeostasis (3–5). Several mutations leading to dysfunction and deregulation of Wnt signaling have been recognized as the etiological agent of multiple growth-related diseases and cancers (6).

The cytoplasmic activation of both canonical (β-catenin dependent) and non-canonical (β-catenin independent) Wnt signaling branches is initiated by the recruitment of Dishevelled protein to the plasma membrane (7). Mammals possess three highly related (8) and mostly functional redundant (9) isoforms of Dishevelled (DVL1-3 in humans). Each DVL protein consists of three structured domains connected by intrinsically disordered regions: N-terminal DIX (Dishevelled and Axin), PDZ (Post-synaptic density protein-95, Disc large tumor suppressor, and Zonula occludens-1), and C-terminal DEP (Dishevelled, Egl-10, and Pleckstrin) (10).

DEP domain can be also found in other protein families where it mostly coordinates the spatial and temporal regulation of the signal transduction by targeting the DEP domain-containing protein to the plasma membrane (11). In Wnt signaling pathway, the membrane targeting of DVL is accomplished via the interaction of DEP domain with Frizzled receptor (12, 13) and negatively charged lipids, in particular phosphatidic acid (PA) (14–17).

The binding to PA membranes occurs via a positively charged patch formed by Helix 3 (residues 462-REARKYASNLLKAG-475) and a contiguous flexible loop (residues 482-NKITFSEQ-489) (16, 17). Interestingly, Hanáková et al. (18) identified four phosphorylation sites at DEP domain of human DVL3 (pS445, pT459, pS469, and pT485), which could perturb the protein-membrane association by, for example, reverting the electrostatic potential (19, 20) or inducing structural modifications (21, 22).

In this study, we combined coarse-grained Monte Carlo (MC) and all-atom Molecular Dynamics (MD) simulations of DEP domain together with experiments of model peptides using quartz crystal microbalance with dissipation (QCM-D). Our multiscale approach allowed us to investigate the structural stability and the binding to PA membranes of phosphorylated DEP in systems with varying lipid compositions, ions in solution, and protonation states of the phosphate groups.

## METHODS

### DEP Domain Structure

The conformation of monomeric DEP domain was based on Dvl2 crystal structure from Yu and coworkers (PDB ID 3ML6) (23). The few necessary mutations to model DEP from DVL3, were performed manually (using PyMOL (24)) as all three human DVL isoforms have high sequence identity with the mouse homolog (Figure S1 and Table S1). The phosphorylated constructs were similarly generated. A comprehensive view of the investigated DEP models can be found in Table 1.

**Table 1:**
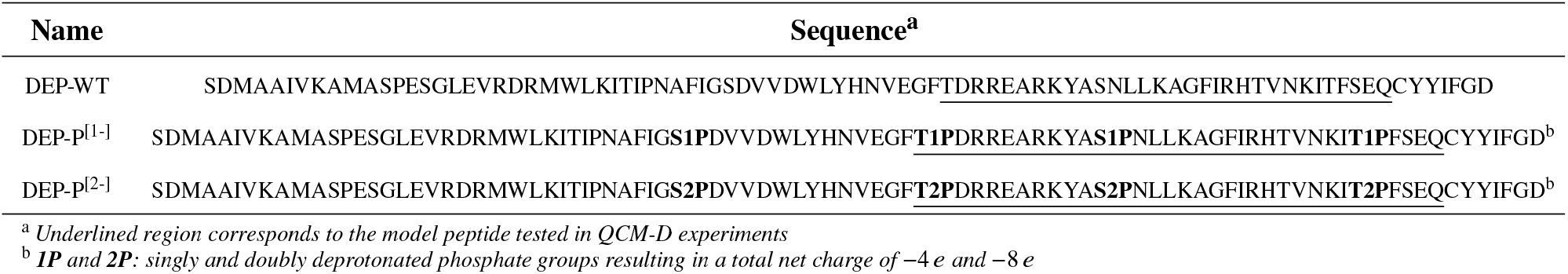
DEP constructs.

### APBS Calculation

The calculations of the surface electrostatic potentials were conducted using the PDB2PQR server (25) and the Adaptive Poisson-Boltzmann Solver (26). For the phosphorylated DEP domain, we extended the AMBER charge scheme (27) with compatible partial charges for phosphorylated residues (28).

### FAUNUS

The interaction between DEP domain and negatively charged membranes was investigated with FAUNUS (29, 30) coarse-grained model and Monte Carlo method. In this model, every amino acid was represented by a single spherical bead, while the membrane surface was modelled by a layer of one hundred beads mimicking phosphatidic acid headgroups. DEP domain was placed above the membrane in the cubic simulation box with dimensions 10 x 10 x 10 nm. Periodic boundary conditions were applied in the XY-plane (i.e. membrane plane). The DEP domain was allowed to freely translate and rotate as a rigid body, while lipid beads could only move in the XY-plane.

The electrostatic energy was computed using either the Debye-Hückel or Coulomb potential, depending on whether the ions were considered implicitly or explicitly. In both cases, the implicit water solvent was accounted for by a relative dielectric constant of 78. In addition, all the species in the system, which can protonate/deprotonate, were subject to titration moves, thus providing an accurate description of the electrostatics at physiological conditions (i.e. pH 7) (29, 30). Table S2 provides an overview of all the investigated systems, including compositions and simulation settings.

To quantitatively evaluate the membrane binding ability of DEP, we calculated the histogram (population) of the distances between the center of mass of DEP and the lipid layer using bin size of 0.1 nm. The associated free energy profile was then computed using Boltzmann inversion.

### All-atom MD Simulations

#### Force Field Parameters

The force field parameters for DEP domain were based on Amber ff99SB-ILDN (31), extended with a compatible set of parameters for phosphorylated amino acids (28). For POPC (1-palmitoyl-2-oleoylsn-glycero-3-phosphocholine) lipid we used Slipids (32–34) parameters, while for singly and doubly deprotonated POPA (1-palmitoyl-2-oleoyl-sn-glycero-3-phosphate) we built a parametrization based on Slipids structural blocks (32–34). A complete description of the parametrization procedure is provided in our previous study of DEP-membrane interaction (17).

Along with the standard non-polarizable force fields, we also employed parametrization with electronic continuum correction (ECC), which accounts for the electronic polarizability (35) via rescaling the partial atomic charges by a factor of 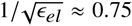, and was shown to significantly improve the interactions of ionic groups (36–41). We extended the ECC scheme from standard charged amino acids to phosphorylated ones, by scaling the charges of the side chain and phosphate group atoms (Table S3). The same scaling was applied to the atoms from the polar regions of singly deprotonated POPA (i.e. headgroup, glycerol backbone, and carbonyl groups). To counteract the reduced hydration of the headgroup, we reduced the s parameter of the Lennard-Jones potential for all the modified atoms following a previously developed ECC model for POPC (42). The final POPA parameters with scaled charges are displayed in Table S4 and a comparison of the membrane properties of the equilibrated lipid bilayers may be found in Figure S2.

### Simulated Systems

#### DEP Domain in Solution

DEP domain constructs based on PDB ID 3ML6 (see Table 1) were solvated with TIP3P (43) water molecules in a cubic box with dimensions of approximately 7 x 7 x 7 nm guaranteeing a distance of at least 1 nm between the protein and the box boundaries. NaCl ions were added at ~150 mM concentration with excess ions to neutralize the system charge.

#### Lipid Bilayers

We studied two lipid compositions: full POPA and POPA:POPC at equimolar concentration. For POPA lipids, two protonation states were considered: singly deprotonated for the pure membrane and exclusively singly or doubly deprotonated for the mixed membrane. The initial configurations, consisting of 64 lipids per leaflet, were generated with CHARMM-GUI (44–46). All systems were then solvated with TIP3P (43) water and counterions were added for electroneutralization. CHARMM-GUI equilibration protocol was run for at least 100 ns at 310 K and 1 bar. Semiisotropic pressure coupling was employed to allow independent changes of the system size in the xy-plane (membrane plane) and z dimensions.

To validate the built POPA parameters, with both standard and ECC charge scheme, we compared the structural properties of analogous systems equilibrated using CHARMM36 (47), apart from doubly deprotonated POPA for which no parameters were available (Figure S2).

#### DEP-Membrane Systems

One DEP domain was positioned above and one below the equilibrated lipid bilayer at approximately 1.5 nm from its surface, with the previously identified PA binding interface (16, 17) oriented towards the membrane. The same solvation as described above for DEP domain in solution was applied. After the first three NPT equilibration steps (see *Simulation Settings*), the center-of-mass of both copies of DEP domain were pulled towards the bilayer along the z-axis over 9 ns, using a force constant of 1000 kJ mol^−1^ nm^−2^. To prevent structural deformations of DEP, the pull rate was very low (0.25 nm ns^−1^). After the pull, the unrestrained equilibration step was performed. Besides NaCl, we also investigated the effect of 150 mM KCl and 10 mM CaCl_2_ when using the ECC charges. The selected model for K^+^ was shown to reproduce the salt activity coefficient derivatives as well as the Kirkwood-Buff (KB) integrals (48), and improved short-range ion-ion interaction (38). The compositions of the simulated systems are summarized in Table S5.

### Simulation Settings for DEP in Solution and DEP-Membrane Systems

All the simulations and analysis were performed using GROMACS Molecular Dynamics package (49) and PLUMED library (50, 51). The systems were energy-minimized using the steepest descent algorithm, first with and then without, position restraints with a force constant of 1000 kJ mol^−1^ nm^−2^ in each spatial dimension on all protein heavy atoms and all lipid atoms (when present). A convergence criteria of 500 or 1000 kJ mol^−1^ nm^−1^ was used.

The subsequent equilibration was divided into five steps, for a total of 9 and 12 ns for DEP in solution and DEP-membrane systems, respectively. First, we heated the systems (i.e. NVT ensemble) to the designated temperatures using the velocity-rescaling thermostat (52) with coupling time of 0.1 ps and position restraints applied only on protein heavy atoms (i.e. no position restraints on lipids). Then, four NPT ensemble steps were performed using the Parrinello-Rahman barostat (53) with coupling time of 2 and 5 ps, for solution and protein-membrane systems. During the NPT equilibration steps the position restraints were sequentially removed from all side-chain, backbone (except Cα), and Cα atoms. No position restraints were impose in the last step.

The production runs for DEP in solution were performed in the NPT ensemble at 300 K and 1 bar for 1000 ns using the same thermostat and barostat as in the equilibration with coupling time of 1 and 10 ps, respectively. Identical conditions were applied to 500 ns long runs of the DEP-membrane systems, except for a temperature increase to 310 K to ensure the liquid phase of lipids. The geometry of water and all covalent bonds, were constrained by SETTLE (54) and LINCS (55) algorithms allowing an integration time step of 2 fs.

Long-range electrostatic interactions were treated with the particle mesh Ewald method (PME) (56), using a grid spacing of 0.12 nm, real space cutoff of 1.2 nm, and PME order 4. Short-range Lennard-Jones interactions were shifted to zero at cutoff distance of 1.2 nm. Long-range corrections for energy and pressure were used for the dispersion interactions. Periodic boundary conditions were applied in all directions.

### Analysis of DEP-Membrane Binding

The contribution to the membrane binding of DEP domain structural components (including the phosphorylation sites), was evaluated via the following per-residue descriptors: (1) the average per-frame number of contacts, where any residue-lipid atom pair with a distance lower than 0.3 nm defines a contact, (2) the total number of unique hydrogen bonds existing between each residue-lipid pair, and (3) the residue-membrane interaction energy. In addition, for each protein residue the distance between its center-of-mass (COM) and the COM of the membrane, along the direction parallel to the membrane normal (i.e. z-axis), was used to estimate the insertion depth into the bilayer.

### Quartz Crystal Microbalance with Dissipation (QCM-D) Experiments

A model peptide encompassing Helix 3 and the contiguous flexible loop of DEP domain was used in QCM-D experiments to test the effect of phosphorylation on the binding to 1,2-Dioleoyl-sn-glycero-3-phosphocholine (DOPC) and 1,2-Dioleoyl-sn-glycero-3-phosphatidic acid (DOPA) supported lipid bilayers (SLBs). The peptide contains three out of four phosphorylation sites (see Table 1). Both wildtype and phosphorylated versions of peptide were purchased from CASLO. The presence of several adjacent phosphorylation sites made the synthesis and purification of the latter version particularly difficult, resulting in a sample of lower purity (~70%).

All QCM-D measurements were performed on device QSense Analyzer (Biolin Scientific, Sweden) at a fixed flow rate of 50 μL min. Firstly, we created films of the desired composition in round bottom tubes by drying aliquots of the lipid stock solutions (Avanti, USA) with a gentle stream of air and 4 hours of vacuum desiccation. The lipid films were then dissolved in ethanol (Avantor, USA) to a final concentration of 0.2 mg mL^−1^. Subsequently, the lipid/ethanol solutions, together with the peptide buffer (PBS and 150 mM at pH 7.4), were used to create the SLBs on SiO_2_ sensors (QSX 303, Biolin Scientific, Sweden) via the solvent-assisted lipid bilayer formation method (57). Additional conditions at higher pH (PBS and 150 mM NaCl at pH 9) or in the presence of Ca^2+^ ions were also tested. To avoid the formation of calcium precipitates in PBS, we employed a two-step approach. SLBs formed in the presence of 150 mM NaCl and 300 mM CaCl_2_ were first washed with 150 mM NaCl alone and then equilibrated with PBS and 150 mM NaCl. In all conditions, stable SLBs had to be observed for at least 5 minutes prior to the injection of the peptide. Each measurement was done at least in triplicate.

## RESULTS

### Impact of Phosphorylation on the Surface Electrostatic Potential of DEP Domain

To evaluate the impact of phosphorylation, the surface electrostatic potentials of both DEP-WT and DEP-P^[2−]^ (phosphorylated DEP with doubly deprotonated phosphate groups) were calculated using the Poisson-Boltzmann approximation (see Figure 1). As expected, DEP-WT features a clear and uniform region of positive potential on the membrane binding side, composed of amino acids from Helix 3 (residues 462-475) and a contiguous loop (residues 482-489) (14, 16). This positive patch was shown to be responsible for the strong electrostatic attraction of the domain to negatively charged lipids, in particular PA (15–17). When residues S445, T459, S469, and T485 are phosphorylated, the electrostatic potential displays a more irregular pattern, with a large negative region occupying almost the whole membrane binding interface. No significant changes occur on the rest of the protein (opposite side from membrane), where the potential only became mildly more negative after phosphorylation (see Figure S3). Note that we used a charge of −2 *e* for the phosphorylated residues, because the phosphate group is mostly doubly deprotonated at physiological conditions (i.e. pH 7) (58–61).

**Figure 1:**
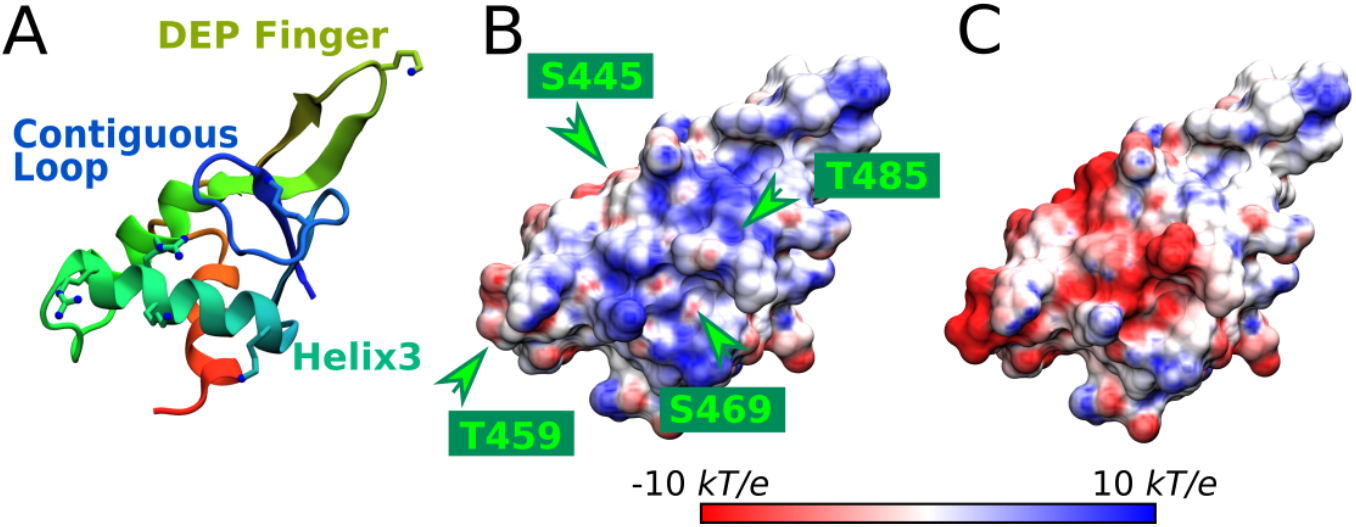
(A): Cartoon representation of DEP domain highlighting the key structural elements. The side chain of the residues forming a polybasic motif are shown as balls and sticks. (B) and (C): Surface electrostatic potential for DEP-WT and DEP-P^[2−]^. The green arrowheads indicate the phosphorylation sites.

### Coarse-grained Simulations

The hypothesis that phosphorylation decreases the affinity of DEP domain to anionic membranes via the perturbation of the electrostatic interaction, was tested using FAUNUS (29, 30) coarse-grained model (see Methods). This model enables to treat ions in solution either implicitly or explicitly and sample the protonation state of both amino acids and lipids.

Figure 2 shows the binding free energy profiles of DEPWT and DEP-P at PA membrane. In implicit salt, DEP-WT was attracted to the membrane with a clear free energy minimum of −5 kJ mol^−1^ at 2 nm distance between DEP center of mass (COM) and the membrane layer. In contrast, DEP-P was repelled, with a shallow unstable minimum at approximately 3 nm. The change from attraction to repulsion was anticipated based on electrostatics because upon phosphorylation the overall charge of DEP domain changed from roughly 0.5 *e* to −6.2 *e*, reflecting a charge per phosphoresidue in the range of −1.5 *e* to −2.0 *e*. However, implicit salt cannot capture the fluctuations of the local ion concentration or ion mediated interactions. Therefore, we also simulated the systems with explicit salt. The obtained free energy profiles are displayed in Figure 2B. DEP-WT adsorbed to the membrane at a distance comparable to the implicit ion model but stronger (−6.2 kJ mol^−1^). Interestingly, DEP-P was also attracted to the membrane with a shallow (−1 kJ mol^−1^) global free energy minimum at the distance of 2.6 nm. The radial distribution function analysis showed that the explicit ions formed a shell around the protein and effectively screened the electrostatic repulsion (Figure S4).

**Figure 2:**
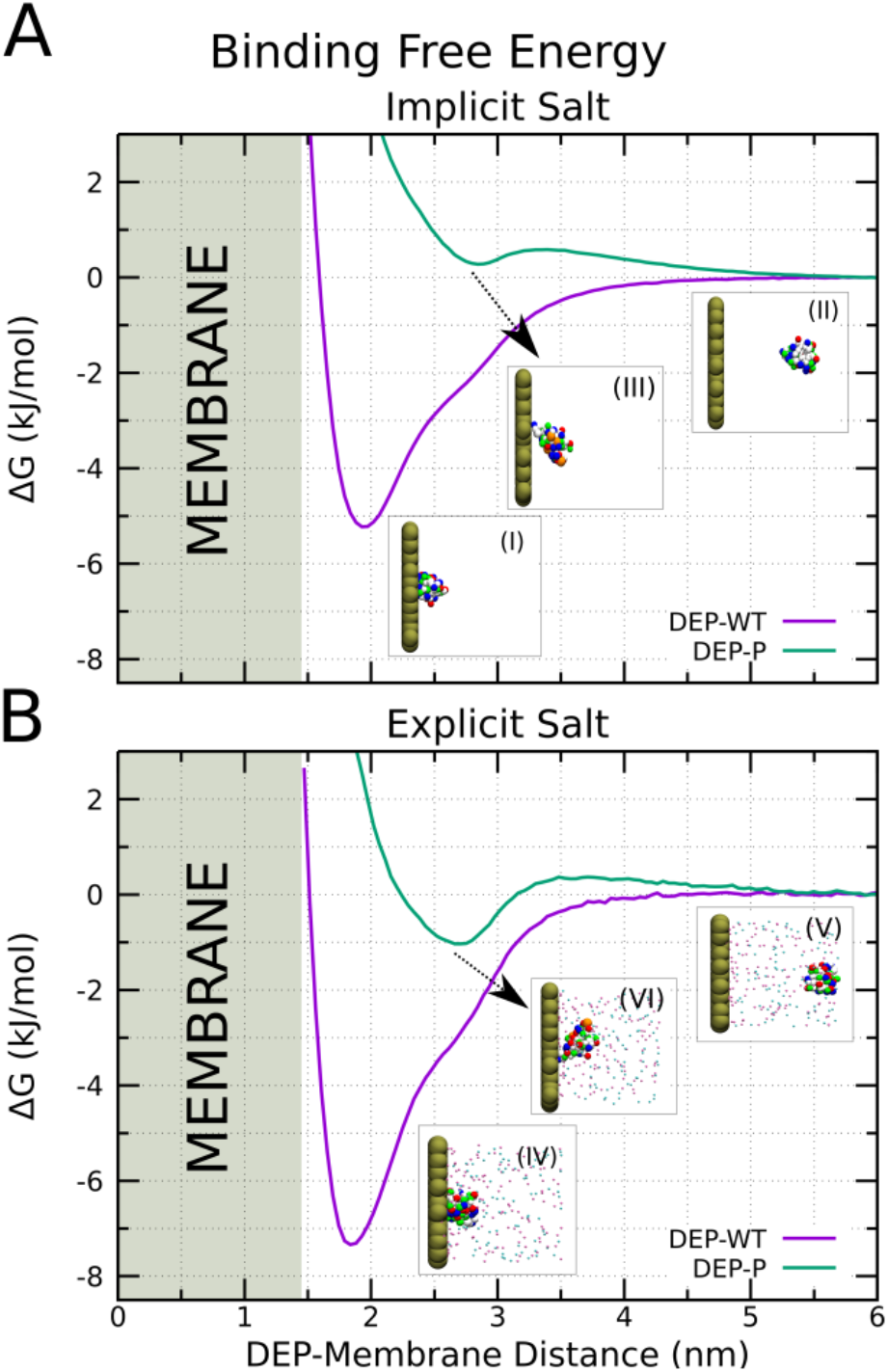
Free energy profiles of DEP domain adsorption to PA membrane in implicit (A) and explicit (B) salt, computed using FAUNUS coarse-grained model. The insets show representative bound and unbound conformations. PA headgroups, Na^+^, and Cl^−^ are shown as tan, pink, and cyan beads. Polar, apolar, basic, acidic, and phosphorylated amino acids are green, white, blue, red, and orange, respectively.

The unexpected binding of negatively charged DEP-P to also negatively charged PA membrane was in different configuration than DEP-WT. The per-residue distributions of the distances in the binding minima (Figure 2 (I) and (IV)), revealed that the binding interface of DEP-WT is mainly composed by a stretch of positive and polar residues belonging to Helix 3 (residues 462-REARKYASNLLKAG-475), a contiguous loop (residues 482-NKITFSEQ-489), and the tip of the DEP finger (residues 434-LKI-436), regardless of the used salt model (see Table S6). In contrast, the free energy minima of DEP-P are mainly populated with conformations showing the DEP finger oriented towards the membrane, while the phosphosites were oriented away from it (Figure 2 (III) and (VI)). We also note that, in general, the presence of explicit ions resulted in a closer binding of the domain compared to the implicit treatment.

### All-atom Simulations

To verify the results from the coarse-grained model and obtain more insight into the role of ions in DEP-membrane interaction, we performed all-atom molecular dynamics simulations.

Firstly, we investigated the domain stability in acqueous solution and compared the per-residue Root Mean Square Fluctuations (RMSF) of all DEP domain constructs. In 1 μs long simulations, DEP-WT and DEP-P^[1−]^ displayed a similar behavior with no significant structural changes (see Figure 3 A and B, respectively). In contrast, the doubly deprotonated phosphorylated residues in DEP-P^[2−]^ caused substantial instability in the DEP finger region (Figure 3 C). Nevertheless, the hydrophobic core formed by the helix bundle was preserved unaltered and the overall fold of the domain remained stable in all cases.

**Figure 3:**
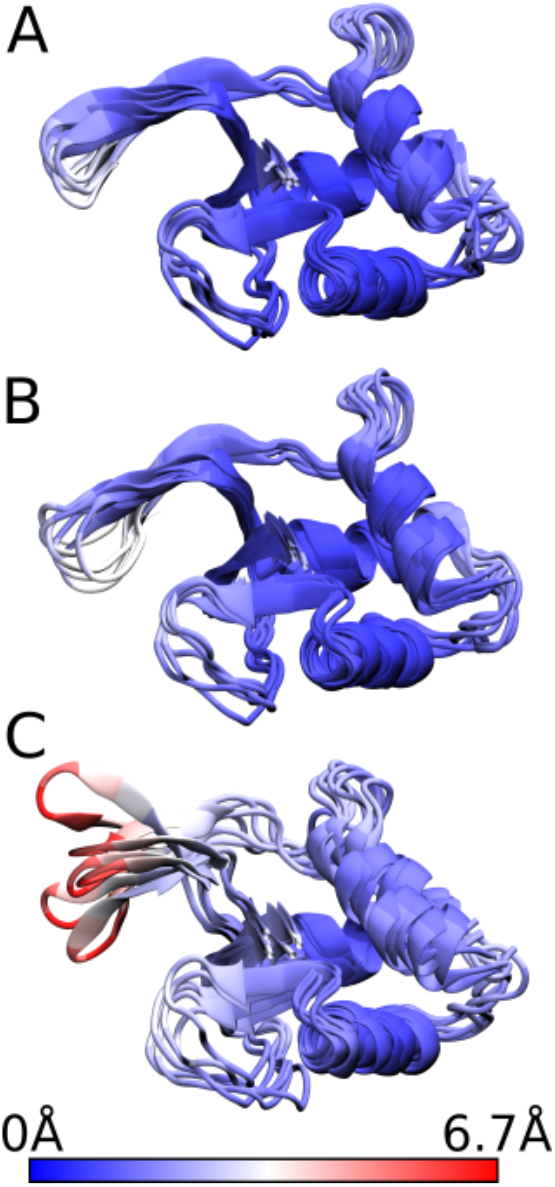
Superimposed conformations from 1 μs long all-atom MD simulations of DEP-WT (A) and DEP-P[1−] (B), and DEP-P[2−] (C). The structures are colored based on the Root Mean Square Fluctuation (RMSF).

The phosphate groups of the doubly deprotonated phosphoresidues were involved in specific interactions with the surrounding local environment (see Table S7 and Figure S16). For example, pS469 formed a strong intramolecular salt-bridge with the neighboring basic amino acid K473. pT485 was also engaged in intramolecular interactions, but the salt-bridge with K483 had lower stability compared to the one of pS469. pT459 phosphate group was mostly interacting with the positive ions in solution. An intermediate behavior, characterized by interactions with other DEP residues and the cations, was shown by pS445. The broad spectrum of local interactions suggests that each phosphorylation modification could underlie a distinct regulatory mechanism as further discussed in the section *Biological Framework*.

After confirmation of the domain stability in solution, we investigated DEP-membrane interaction. The upper panels of Figure 4 show the final snapshots of 500 ns long simulations of DEP-WT (A), DEP-P^[1−]^ (B), DEP-P^[2−]^ (C) at POPA membrane. In all cases, the domain adopted a similar bound conformation, with Helix 3 (residues 462-475) and a contiguous loop (residues 482-489) in steady contact with the lipid bilayer, in agreement with previous simulations of DEP-WT (16, 17). The residues at the tip of the DEP finger (residues 434-436) also contributed to the binding, but the flexibility of the loop resulted in less stable interactions with the membrane (Figure S5). Two phosphorylation sites (pS469 and pT485) right in the middle of the membrane binding interface, and a third one (pT459) located on the side, were regularly in contact with the membrane surface (Figure 4 B and C). Despite their electrostatic repulsion, the phosphate groups on the protein were in the vicinity of POPA headgroups and contributed to the binding via a network of ion-mediated interactions, where a cation interacted with two negatively charged oxygens from different phosphate groups (see insets of Figure 5). This behavior was particularly enhanced for the doubly deprotonated phosphoresidues. To investigate deeper the nature of the ion-mediated interaction and ensure that the results are not model and system dependent, we employed: (1) a model with scaled charges (ECC), (2) different salt cations, (3) a lipid mixture with zwitterionic POPC lipids, and (4) fully deprotonated POPA lipids.

**Figure 4:**
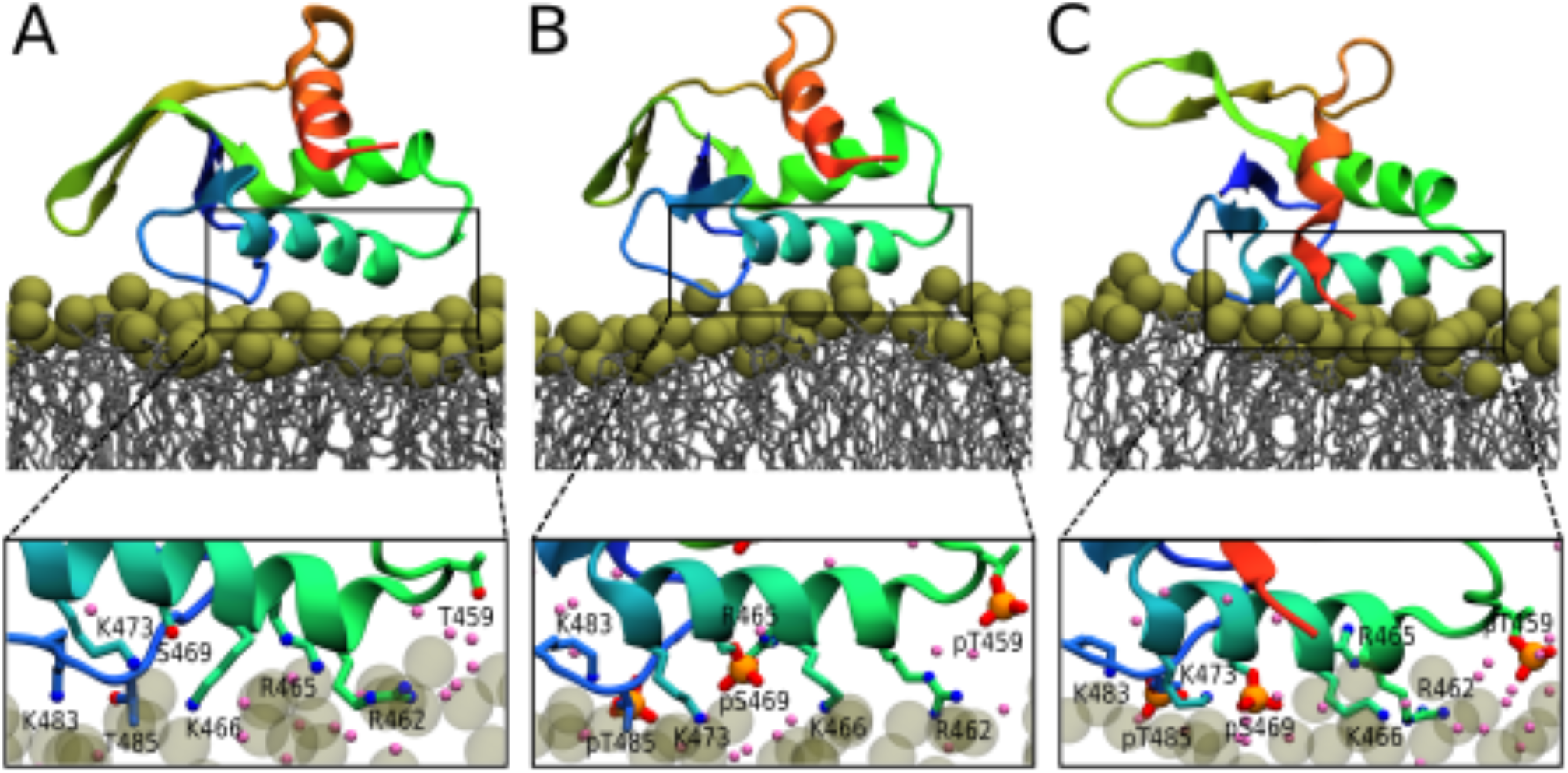
Final snapshots from 500 ns long all-atom simulations of DEP-WT (A), DEP-P^[1−]^ (B), and DEP-P^[2−]^ (C) in the presence of POPA membrane. Upper panels: overview of the bound configurations. The mild rotation of the domain in panel C is indicative of the dynamics nature of the interaction with the membrane, during which Helix 3 (residues 462-475) and the contiguous loop (residues 482-489) remained steadily bound. Lower panels: close-ups of the binding interface highlighting the critical residues. DEP domain is displayed in cartoon representation. Tan spheres represent POPA headgroup, while the glycerol backbone and the acyl tails are shown as gray sticks. Red, blue, and pink spheres are for O, N and Na^+^ atoms, respectively. P and O atoms in the phosphate group of the phosphorylated residues are displayed as orange VDW sphere and red sticks. Cl^−^ ions and water molecules are omitted for clarity.

**Figure 5.**
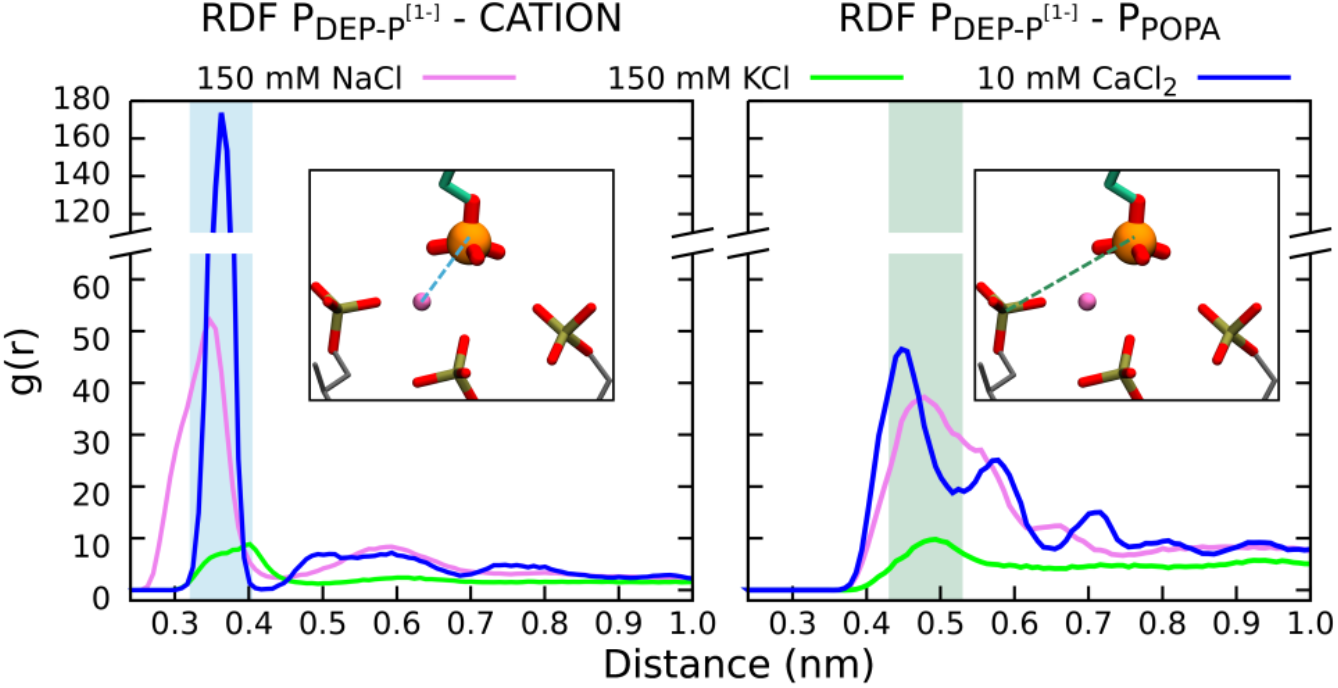
Radial Distribution Function (RDF) between P atoms from DEP-P^[1−]^ phosphorylated residues and cations (left panel) or P atoms from POPA headgroups (right panel). The representative configurations of the corresponding atom pairs shown in the insets illustrate the ion-mediated interactions between the phosphate groups from POPA lipids and DEP domain phosphoresidues. Note the narrow distribution (0.1 nm) of the positions of the highest peaks in both plots (shaded areas). All systems were simulated with ECC charges. The color scheme of the insets is the same as in Figure 4.

### (1) Scaled Charges

To prevent the overestimate of the ion binding reported for common non-polarizable force fields (36, 37, 62), we applied the electronic continuum correction (ECC) method (35, 42, 63), which effectively includes the contribution of the electronic polarizability.

The ECC model significantly decreased the accumulation of Na^+^ at the lipid interface (Figure S6), in agreement with previous observations for POPC or POPS membrane (42, 63, 64). Despite the reduced electrostatic interactions between charged species, the protein regions required for the binding of DEP-P^[1−]^ to POPA membrane, were the same as in the case of the non-scaled model (Figure S7). Moreover, the broader density peak for the phosphate headgroup exhibited by the POPA-ECC model, possibly indicates a looser packing of the lipid molecules, resulting in deeper membrane insertion of the phosphate groups of DEP-P^[1−]^ (Figures S7 and S8) and domain coordinated Na^+^ ions (Figure S6).

### (2) Ion Types

The effect of different ions on DEP domain binding to anionic membranes was evaluated by performing additional simulations using the ECC scheme and solutions with ~150 mM KCl and ~10 mM CaCl_2_. K^+^ was chosen as the most abundant cation in the cytoplasm, while Ca^2+^ represents a well-known second messenger, capable to interact with lipid membranes and proteins.

In general, the same structural regions of DEP-P^[1−]^ were engaged with the membrane when replacing Na^+^ with K^+^ or Ca^2+^ (Figure S7). However, the ions displayed different affinity for the lipid headgroups, with Ca^2+^ featuring the strongest interaction, followed by Na^+^ and K^+^ (Figure S9). Figure 5 shows the radial distribution function (RDF) between the P atom from DEP phosphorylated residues and system cations or the P atoms of POPA headgroups. The first peak in RDF corresponds to the closest preferred contact and its height correlates with the strength of the attraction. Phosphorylated residues thus made similar close contacts with all cations, with preference Ca^2+^ » Na^+^ » K^+^. We conclude that all investigated ions could mediate the interaction between the phosphate groups of DEP-P^[1−]^ and POPA lipids, although the stability of such interaction was dramatically affected by the nature of the ion.

### (3) Membrane composition

In mammalian cells, PA accounts for a small portion of the total phospholipid content (i.e. ~1%) (65, 66). Even though PA local concentration can be increased by metabolic biosynthesis (67, 68) and microdomain segregation (69, 70), a lipid patch composed of only POPA might represent an extreme scenario. Therefore, we also investigated a membrane with POPA and POPC lipids in a 1:1 ratio.

We observed that the membrane binding ability and the binding sites of DEP-WT, DEP-P^[1−]^, and DEP-P^[2−]^ were not significantly affected (Figures S5 and S10), suggesting that even a lower local enrichment of PA could be sufficient for a stable recruitment of the domain. To inspect the local surrounding of DEP, we calculated the radial distribution function (RDF) of P atom from POPA or POPC lipids with respect to the protein surface (Figure 6). Irrespective of the phosphorylation state, DEP domain preferentially interacted with PA lipids and, on average, approximately five PA molecules were located within 0.3 nm of the domain, compared to only one PC molecule (see insets in Figure 6). Consistently, in any given time of the simulations, DEP domain and the phosphorylated residues involved in ion-mediated interactions (pT459, pS469, and pT485) participated in roughly four times (or higher) more hydrogen bonds with PA than PC lipids (Table S8).

**Figure 6:**
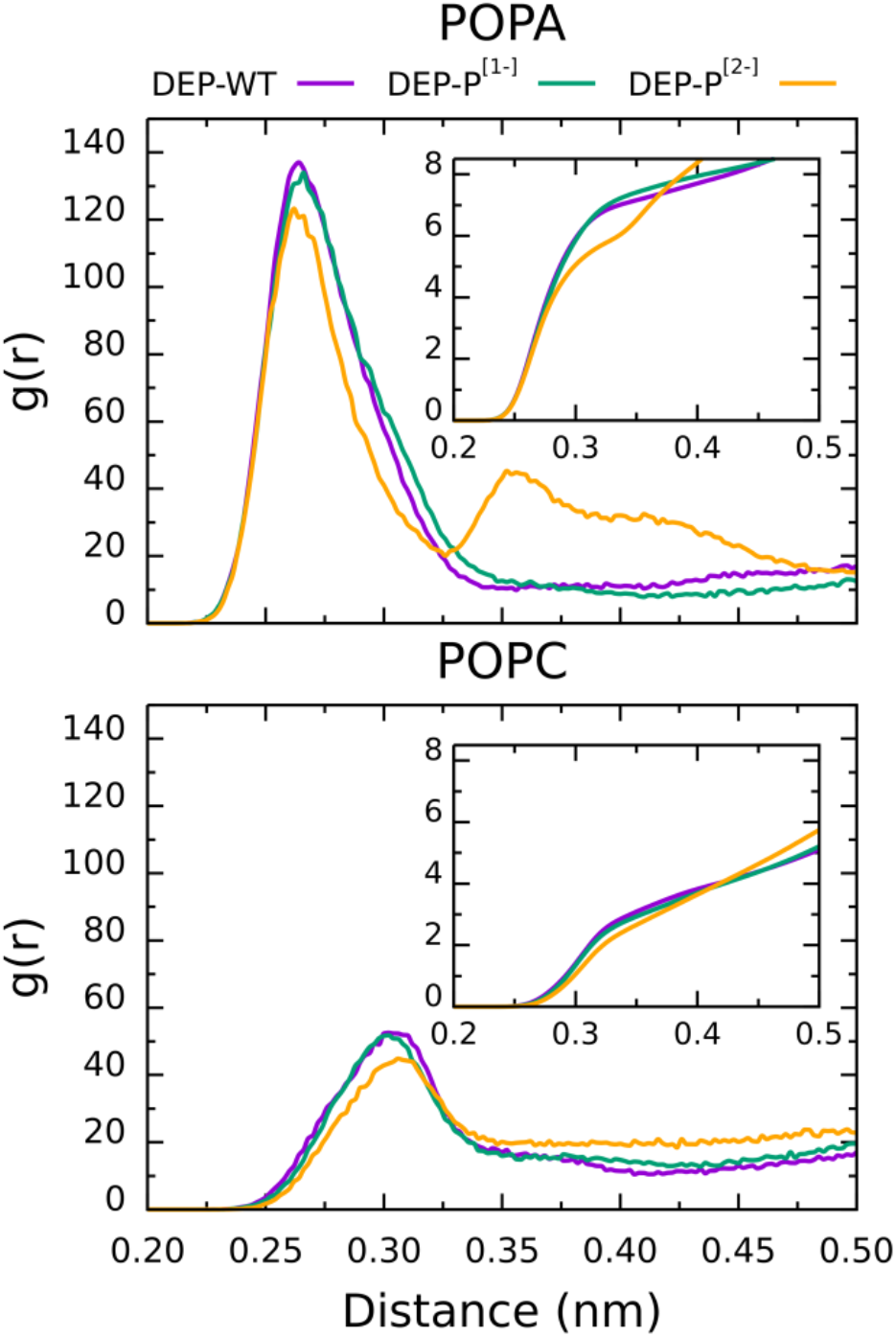
Radial distribution function (RDF) of P atom from POPA (upper panel) and POPC (lower panel) lipids around DEP domain surface. The cumulative number is displayed in the insets. The profiles were averaged over both lipid leaflets and DEP domain copies.

Note that in Figure 6, DEP-P^[2−]^ shows a second peak in the RDF profile for PA at ~0.35 nm. Given that the lipid aggregation does not significantly differ in the investigated systems (Figure S11), it is possible that the higher cations coordination number of DEP phosphoresidues (Table S7) led to more branched ion-mediated interactions with PA headgroups (Figure S12).

### (4) PA Protonation State

The phosphomonoester headgroup of PA is characterized by two pK_a_, one of which has a value in the physiological pH range (71). As a consequence, minor changes in the local environment could lead to full deprotonation. Therefore, in the mixed POPC:POPA bilayer, we also used a fully deprotonated model of POPA with charge −2 *e* (see Methods). As expected, the loss of the hydrogen bond donor dramatically reduced the tendency of PA to form hydrogen bonds with DEP domain. However, a stronger electrostatic attraction with positively charged amino acids and ions, resulted in firm interactions with DEP-WT and phosphorylated DEPs, respectively (Figure S13).

The introduction of fully deprotonated PA had also a pronounced impact on the membrane organization with the formation of stable PA aggregates, which are kept together via ion-mediated interactions (Figure S14). Indeed, RDF analysis demonstrated that Na^+^ interacted six times more with dianionic PA than its monoanionic form (Figure S15).

### Phosphorylated Residues

When comparing simulations of DEP-P^[2−]^ in solution and at anionic membranes, we could observe significant changes in the local environment of specific phoshoresidues. For instance, the intramolecular salt bridge formed in solution by pS469 was partially disrupted at full PA membrane, while the overall stability of the weaker pT485 intramolecular interaction was increased at both investigated PA concentrations (Figure S16). Interestingly, the latter event was associated with a local enrichment of positive ions around the phosphate group of pT485 (Table S7), suggesting a possible role in the formation of ion-mediated interactions. In contrast, the local environment of the remaining two phosphoresidues (pT459 and pS445) exhibited no noticeable changes.

### Binding of Helix 3 Model Peptides to Supported Lipid Bilayers

QCM-D measurements provide changes in crystal resonant frequency (Δ*f*) and dissipation (Δ*D*), which are related to the adsorbed mass and its viscoelastic properties. Once the base line is established for SLB, the frequency decrease is related to adsorption of the peptide to the membrane.

Figure 7 shows QCM-D measurements of both model peptides at pure DOPC and DOPA supported lipid bilayers. After the injection of the non-phosphorylated peptide (panel A), we observed strong adsorption to DOPA membrane, together with a residual interaction with DOPC lipids. For the phosphorylated Helix 3 peptide (panel B), in all tested conditions, the magnitude of the signal was generally weaker, suggesting reduced, although non-negligible, affinity. Unlike the unmodified counterpart, the binding to DOPA and DOPC SLBs could not be clearly discriminated. However, the introduction of Ca^2+^ ions significantly strengthened the interaction with DOPA lipids, in good agreement with the results from our simulations shown in Figure 5.

**Figure 7:**
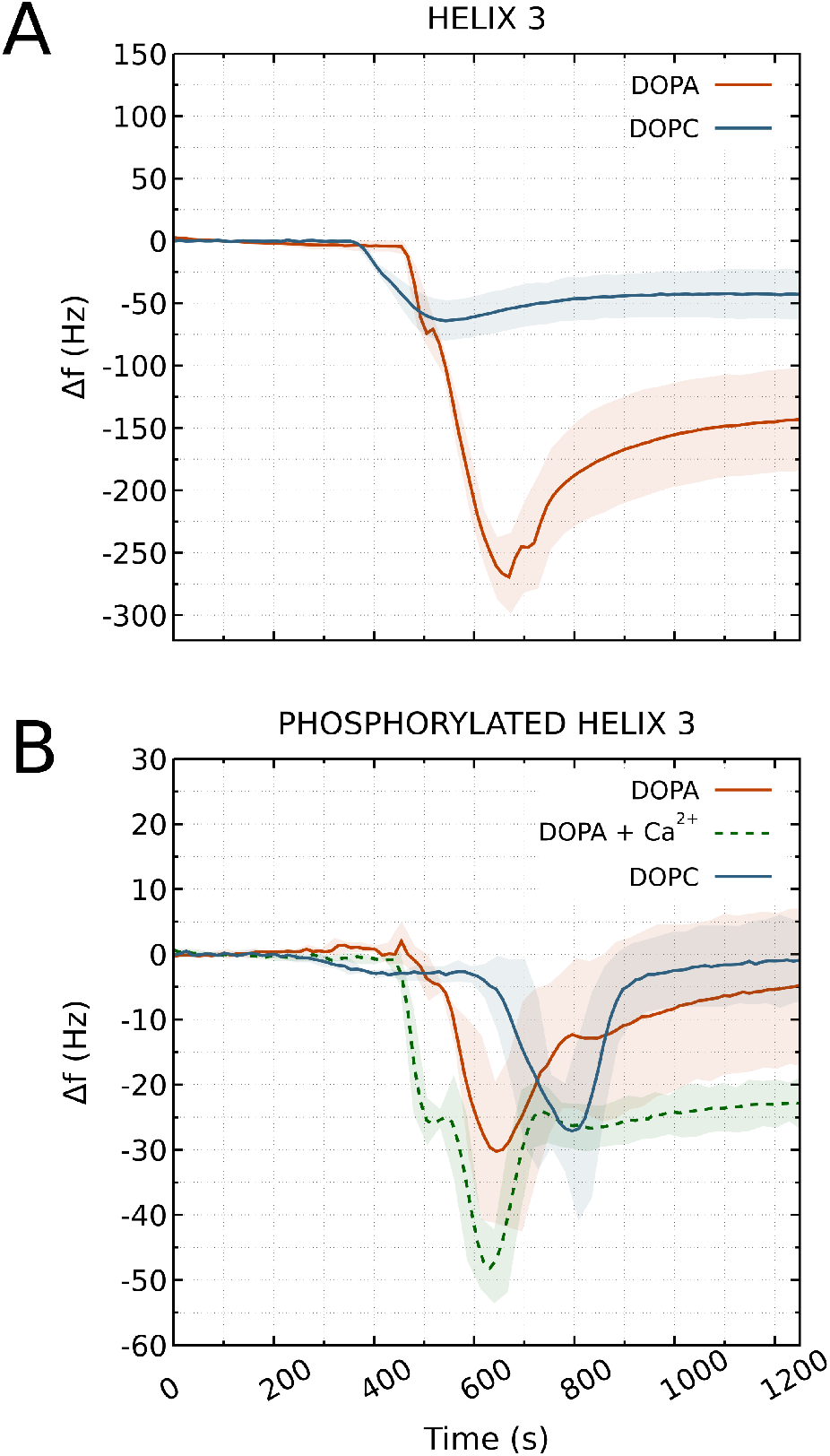
QCM-D experiments of Helix 3 peptide (and its phosphorylated form) at DOPC and DOPA supported lipid bilayers. The extent of the negative frequency change with respect to the stable lipid bilayer (y=0) is proportional to the adsorbed peptide. Data shown as average and standard deviation (shaded area) over three or four separated measurements. Only the third overtone is considered.

## DISCUSSION

The binding of DEP-WT to PA lipid is mainly driven by electrostatic interactions (16, 17). Thus, phosphorylation could significantly decrease the binding affinity by reversing the electrostatic potential of the domain. Here, we report a more complex scenario, where the intimate interaction of phosphorylated DEP with the local environment resulted in stable binding of PA rich membranes.

### DEP Domain Stability

DEP domain was found to be stable in solution and only locally affected by phosphorylation within 1 μs long all-atom simulations. The largest structural fluctuation was associated with the flexible DEP finger and this behavior did not change when the domain was bound to the membrane, regardless of the lipid composition and solution cations.

Therefore, although it cannot be excluded that phosphorylation can in some measure contribute to DEP conformational change (72), it is unlikely that a mechanism of this sort is involved in the direct regulation of DEP-membrane binding.

### Membrane Organization

Previous estimates for the surface pK_a_ of PA monolayers (73, 74) suggested that the lipid is mostly singly deprotonated at physiological pH, a behavior nicely reproduced by our FAUNUS model (Table S2). However, in all-atom simulations, we adopted both singly and doubly deprotonated states, due to the decrease of the second pK_a_ of PA in binary mixtures with PC lipids (75, 76). The high charge density of doubly deprotonated PA induced strong adsorption of Na^+^ to the membrane, resulting in PA aggregation via cation-mediated interactions (77).

Figure 5 shows that Ca^2+^ can also cause local aggregation of singly deprotonated PA (right panel), a behavior documented for other negatively charged phospholipids (78–80) and typically associated with acyl tails ordering. For both PA charge states, we observed either no change or rather small decrease in the acyl tails order parameters (Figure S2), consistent with the putative stabilizing effect of PA aggregates on nAChR receptor (81).

### DEP Domain Interaction with PA Membranes

#### Simulations

Regardless of the computational model and system composition, DEP-WT interacted with PA-rich membranes via a stretch of positively charged amino acids located on Helix 3 (residues 462-475) and a contiguous flexible loop (residues 482-489) (Figures 4 and S10). Altogether these residues form a polybasic motif, a structural feature frequently observed in many proteins bound to the cytoplasmic leaflet of the plasma membrane (82–85).

The behavior of phosphorylated DEP was strongly influenced by the ion representation and force field used. In coarse-grained simulations with implicit ions, the domain association to the membrane was prevented. Instead, we observed favorable binding of DEP-P finger to the anionic membrane (Figure 2 VI), when the ions were explicitly treated. Interestingly, a secondary binding mode with the phosphoresidues oriented towards the membrane anticipated the presence of ion-mediated interactions (Figure S17), a bound conformation greatly stabilized in all systems simulated at all-atom resolution. Indeed, the phosphorylated residues (pT459, pS469, and pT485) formed both ion-mediated and hydrogen bond interactions, which are not fully captured by the employed coarse-grained model, with PA membranes in all-atom simulations. Notably, the dense hydrogen bonding could promote the full deprotonation of PA headgroup (86), which in turn would result in stronger attraction of the solution cations and enhanced ion-mediated interactions. Similar effect on PA headgroups could be induced by phosphatidylethalonamines (75), the most abundant zwitterionic phospholipids in the inner leaflet of the plasma membrane (87, 88), and multivalent cations (89), such as Ca^2+^. Ca^2+^ is a known secondary messenger in non-canonical Wnt pathways (90) and, in our simulations, was bound between DEP-P phosphorylated residues and PA lipids with stronger affinity than monovalent ions, in line with previous studies on the ion binding to protein charged groups and lipid membranes (39, 91–94). Thus, a scenario where the fine regulation of the local Ca^2+^ concentration mediates the cross-talk between post-translational modifications (i.e. DEP phosphorylation) and membrane binding, represents an intriguing possibility.

#### Experiments

Quantitative and efficient site-specific protein phosphorylation has proven rather challenging in *in vitro* systems and alternative approaches, such as phosphomimicking (mutation of the phosphosites to Asp and Glu), do not fully capture the physico-chemical nature of phosphate groups. Thus, to study how phosphorylation affect DEP-membrane interaction, we decided to use synthetically prepared (phospho-)peptides covering most of the amino acids forming the PA-binding interface and three out of four phosphoresidues.

Measurements of the non-phosphorylated peptide at diunsaturated lipids, chosen for their fluid state at room temperature, showed an expected binding preference for PA (Figure 7 – panel A). The observed residual affinity for PC membrane could originate from attraction between lipids and peptide residues which are missing their binding partners from the domain. Despite the electrostatic repulsion, we detected also a non-negligible binding between the negatively charged peptide and PA lipids (Figure 7 – panel B). In line with our simulations, this interaction was enhanced when Ca^2+^ ions were injected before the peptide.

Our results suggest that phosphorylation of DEP domain does not act as a simple electrostatic switch, as reported previously for other membrane binding proteins (19, 20, 95). In those studies, the examined model membranes mainly contained phosphatidylglycerol, phosphatidylserine, and phosphatidylinositol as anionic species. Moreover, when PA was considered, its molar concentration was ~10%, which is significantly less than in the present work, focused on PA-rich membranes. Therefore, we believe that the cation accumulation at the membrane surface, and consequently the related screening of the electrostatic repulsion and the formation of ion-mediated interactions, were considerably less pronounced than in our systems. We conclude that the effect of phosphorylation on DEP membrane-binding ability could depend not only on the local lipid and salts concentration, but also on the specific nature of the anionic phospholipid (i.e. PA).

### Role of Individual Phosphorylation Sites

In this final section we describe the role of DEP domain individual phosphorylation sites in the context of membrane binding and protein-protein interactions.

pT459 was frequently found in the proximity of the membrane surface, however the contribution to the hydrogen bonding of the lipid molecules was rather negligible (see for example Table S8 for DEP at POPA:POPC membrane). Unlike the other phosphoresidues engaged with the membrane, pT459 is located outside the membrane binding interface, on a solvent exposed and unstructured region connecting Helix 1 and Helix 2 (Figure 1). The observed ion-mediated interactions of this residue with PA lipids thus likely originates from affinity for solution cations rather than a specific membrane binding mechanism, as suggested by the strong coordination of positive ions with both the membrane bound and unbound conformations (Table S7). Therefore, we are unable to draw any firm conclusions on how T459 phosphorylation affects membrane binding.

pS469 is located on Helix 3, where it strongly interacted with the nearby amino acids belonging to the polybasic motif, especially K473. The formed intramolecular salt-bridge could reduce DEP-membrane affinity by sequestering the lysine and preventing its interaction with PA-rich membranes. An inhibitory effect on membrane binding was also reported for phosphorylation of the first DEP domain of P-Rex1 (95). Interestingly, phosphorylation by RIP4K kinase on S480, the equivalent of S469 in DVL2, stimulates the canonical Wnt signalling response by enhancing the cytoplasmic assembly of active DVL2 signalosomes β-catenin (96). Finally, we note that in our simulations, the salt bridge between pS469 and K473 was partially disrupted upon binding to the full PA membrane (Figure S16), suggesting that very high PA concentration might overcompensate for the proposed downregulation of DEP membrane association by increasing the likelihood of forming ion-mediated interactions with pS469.

The flexible loop (residues 482-489) containing pT485 was prone to local structural fluctuations. Therefore, in solution, the salt bridge involving pT485 and K483 was less stable than the one formed by pS469 (Figure S16 – lower panels). As the domain approached the membrane, the salt bridge stabilised, allowing the cations adsorbed on the bilayer surface to interact with the phosphate group of pT485 and anchor DEP-P via ion-mediated interactions. At the membrane surface, pT485 formed direct hydrogen bonds with PA headgroups, thereby strengthening DEP-membrane association (Table S8). No significant correlation between PA content and salt bridge stability was observed, suggesting that pT485 may support ion-mediated interactions over a wide range of PA concentrations.

Anionic lipids are not the only interacting partners of DEP domain. In fact, a surface dipole created by the tip of DEP finger (K435) and residues D446 and D449 was predicted to mediate the interaction with upstream regulators (14), as further confirmed by the impairment of FZD-dependent membrane localization of DVL protein caused by K435M point mutation (12, 97, 98). Consistently, the proximal pS445 did not form any stable direct or ion-mediated interaction with PA lipids. Instead, S445 phosphorylation could be responsible for the higher conformational plasticity exhibited by the DEP finger (Figure 3C), a structural feature that may be linked to homo-dimerisation of the DEP domain (99) and formation of active signalosomes.

## CONCLUSION

Using computational and experimental approaches, we investigated the effect of phosphorylation on the structural stability and membrane binding of DEP domain from human Dishevelled protein. We hypothesized that the electrostatic-driven binding of DEP to negatively charged membranes (16, 17) could be reverted by four recently identified phosphorylation sites (18).

Unexpectedly, the strong adsorption of the cations onto the surface of PA-rich membranes, effectively screened the electrostatic repulsion and enabled ion-mediated and hydrogen bond interactions between the phosphorylated residues of DEP domain and the lipid headgroups. The emerging scenario suggests that DEP-membrane association could depend on the complex interplay between phosphorylation and the local concentration of ions and PA lipids. We believe that our findings help to elucidate the regulation of Wnt signaling pathway and, broadly speaking, other biological processes involving protein-membrane association and post-translational modifications.

## SUPPLEMENTARY MATERIAL

Supplementary material is available online.

## AUTHOR CONTRIBUTIONS

R.V. designed the research. F.L.F. carried out all simulations, analyzed the data. M.D. carried out QCM-D measurements. V.B., R.V., and F.L.F. wrote the article.

## ACKNOWLEDGMENTS

The work was supported by the Czech Science Foundation (grant GA20-20152S) and the project National Institute of virology and bacteriology (Programme EXCELES, ID Project No. LX22NPO5103) – Funded by the European Union – Next Generation EU. Computational resources were provided by the CESNET LM2015042 and the CERIT Scientific Cloud LM2015085 provided under the program Projects of Large Research, Development, and Innovations Infrastructures. Additional computational resources were obtained from IT4 Innovations National Supercomputing Center – LM2015070 project supported by MEYS CR from the Large Infrastructures for Research, Experimental Development and Innovations. We also acknowledge CF Prot of CIISB, Instruct-CZ Centre, supported by MEYS CR (LM2018127) and European Regional Development Fund-Project „UP CIISB“ (No. CZ.02.1.01/0.0/0.0/18_046/0015974)

