## Supplementary Material for "Counterintuitive Binding of Phosphorylated DEP Domain from Dishevelled Protein to Negatively Charged Membranes"

*Francesco L. Falginella, Martina Drabinová, Vítězslav Bryja, and Robert Vácha*

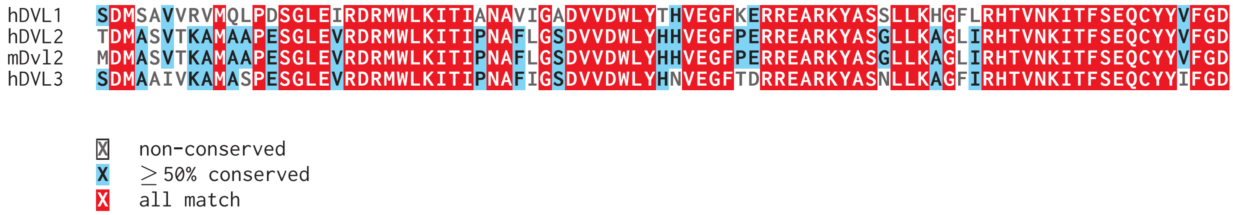


**Figure S1:** Multiple sequence alignment of DEP domain from human (h) DVL1, DVL2, DVL3 and mouse (m) Dvl2. The alignment was generated using Clustal Omega (100)

**Table S1: Sequence identity matrix for human (h) DVL1, DVL2, DVL3 and mouse (m) Dvl2 in percentage.**

| | **hDVL1** | **hDVL2** | **mDvl2** | **hDVL3** |
|---|---|---|---|---|
| **hDVL1** | 100 | 77 | 77 | 78 |
| **hDVL2** | | 100 | 99 | 86 |
| **mDVL2** | | | 100 | 86 |
| **hDVL3** | | | | 100 |

*The alignment was generated using CLUSTAL O(1.2.4)*

**Table S2: FAUNUS Systems**

| Protein[a] | Non-bonded[b] | Salt[c] | Lipid Charge[d] |
|---|---|---|---|
| DEP-WT | Debye-Hückel | Implicit | $-1.17\,e$ |
| DEP-P | Debye-Hückel | Implicit | $-1.17\,e$ |
| DEP-WT | Coulomb | Explicit | $-1.08\,e$ |
| DEP-P | Coulomb | Explicit | $-1.05\,e$ |

[a] *pKa values: 4.0 (D), 4.4 (E), 6.3 (H), 12.0 (R), 10.8 (C), 9.6 (Y), 10.4 (K), 2.19 ($pKa_1$ - pS), 5.78 ($pka_2$ - pS), 6.10 ($pka_2$ - pT), 5.9 ($pka_2$ - pY)*
[b] *Listed electrostatic potential together with repulsive isotropic Lennard-Jones interactions*
[c] 150 mM *NaCl. Explicit salt includes excess ions for electroneutralization*
[d] *Per lipid bead charge averaged over 100 PA lipids ($pKa_2$ = 6.9)*

**Table S3: Extended ECC charge scheme for phosphorylated amino acids**

| **Singly Deprotonated Phosphate Group[a]** | | | | | |
|---|---|---|---|---|---|
| **pSer** | | **pThr** | | **pTyr** | |
| *Atom* | *Charge (e)* | *Atom* | *Charge (e)* | *Atom* | *Charge (e)* |
| CB | 0.015691 | CB | 0.334476 | HD1 | 0.129914 |
| HB1 | 0.095851 | HB | 0.019263 | HE1 | 0.169950 |
| HB2 | 0.095851 | OG1 | -0.368072 | HE2 | 0.169950 |
| OG | -0.349268 | P | 0.946457 | HD2 | 0.129914 |
| P | 0.992226 | O1P | -0.529013 | OG | -0.400839 |
| O1P | -0.564586 | O2P | -0.611516 | P | 1.044910 |
| O2P | -0.616358 | O3P | -0.611516 | O1P | -0.564616 |
| O3P | -0.616358 | H1P | 0.314307 | O2P | -0.616848 |
| H1P | 0.320326 | | | O3P | -0.616848 |
| | | | | H1P | 0.317487 |

| **Doubly Deprotonated Phosphate Group[a]** | | | | | |
|---|---|---|---|---|---|
| **pSer** | | **pThr** | | **pTyr** | |
| *Atom* | *Charge (e)* | *Atom* | *Charge (e)* | *Atom* | *Charge (e)* |
| CB | 0.254462 | CB | 0.604787 | HD1 | 0.112007 |
| HB1 | 0.043887 | OG1 | -0.408923 | HE1 | 0.161111 |
| HB2 | 0.043887 | P | 1.024542 | HE2 | 0.161111 |
| OG | -0.322757 | O1P | -0.724280 | HD2 | 0.112007 |
| P | 1.040315 | O2P | -0.724280 | OG | -0.418458 |
| O1P | -0.737215 | O3P | -0.724280 | P | 1.035504 |
| O2P | -0.737215 | | | O1P | -0.707663 |
| O3P | -0.737215 | | | O2P | -0.707663 |
| | | | | O3P | -0.707663 |

[a] *The original non scaled charges were taken from (28)*

**Table S4: Extended ECC charge scheme for PA lipid**

| Singly Deprotonated PA Headgroup[a] | |
|---|---|
| *Atom* | *Charge (e)* |
| P | 0.732 |
| C1 | -0.075 |
| C2 | 0.353 |
| C3 | 0.150 |
| O11 | -0.319 |
| O12 | -0.480 |
| O13 | -0.589 |
| O14 | -0.589 |
| C21 | 0.593 |
| O21 | -0.353 |
| O22 | -0.488 |
| C31 | 0.593 |
| O31 | -0.353 |
| O32 | -0.488 |
| HA | 0.060 |
| HB | 0.060 |
| HS | 0.053 |
| HY | 0.045 |
| HX | 0.045 |
| H12 | 0.300 |

[a] *For all modified atoms the σ parameter of the Lennard-Jones potential was slightly reduced to account for lipid dehydration.*

**Table S5: DEP-membrane systems investigated with all-atom simulations**

| Lipids | Charge Scheme | DEP[a] | Cation |
|---|---|---|---|
| POPA | Standard | $0\,e$ | $Na^+$ |
| | | $-4\,e$ | |
| | | $-8\,e$ | |
| | ECC | $0\,e$ | $Na^+$,$K^+$,$Ca^{2+}$ |
| | | $-3\,e$ | |
| POPA:POPC[b] | Standard | $0\,e$ | $Na^+$ |
| | | $-4\,e$ | |
| | | $-8\,e$ | |

[a] *Total net charge of DEP-WT, DEP-P[1-], and DEP-P[2-]*
[b] *Singly and doubly deprotonated POPA*

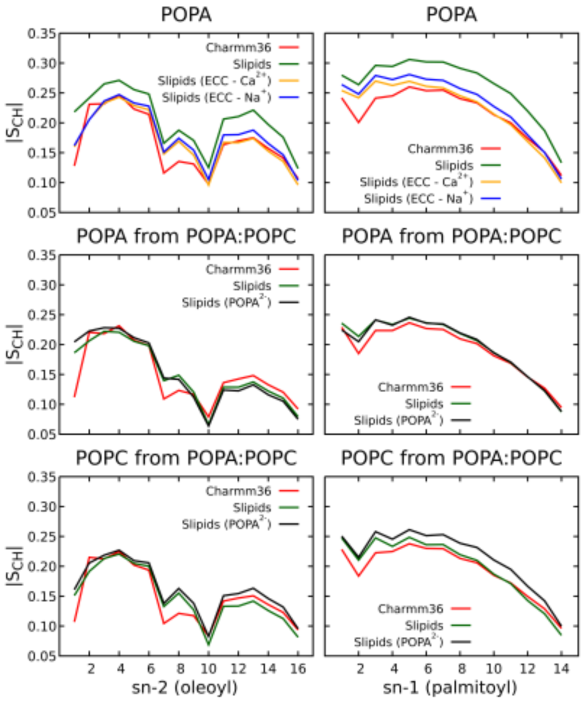


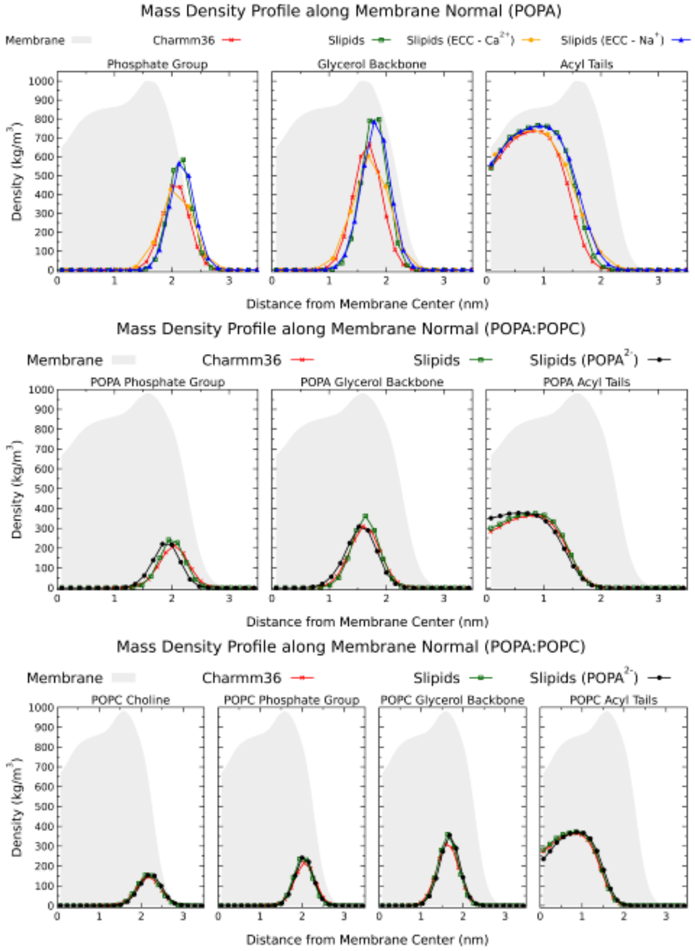


| Force Field | Area per Lipid(Å²) |
|---|---|
| *POPA Membrane* | |
| Charmm36 | 58.7 |
| Slipids | 52.8 |
| Slipids ($Ca^{2+}$)(1) | 52.5 |
| Slipids (ECC) | 53.2 |
| *POPA:POPC Membrane(2)* | |
| Charmm36 | 61.7 |
| Slipids | 59.3 |
| Slipids ($POPA^{2-}$) | 58.1 |

(1) 10 mM $CaCl_2$; (2) Combined Area per Lipid

**Figure S2:** Comparison of the structural properties of the membrane models employed in this study. Top left: Order parameters. The developed PA parameters led to increased ordering compare to CHARMM36, but the resulting area per lipid (bottom left table) is in line with previous estimates (101). Interestingly, the ECC charge scheme increased the bilayer disorder. PA aggregates, induced by either $Ca^{2+}$ or doubly deprotonated PA, show very similar or reduced ordering compared to singly deprotonated PA in the presence of $Na^{+}$ (green line). Note that the data for the bilayer simulated with $Ca^{2+}$ were obtained in the presence of DEP domain, while in the other displayed cases the bilayer was simulated alone. Right panels: Mass density profiles of the lipids building blocks. The gray shaded areas show the whole mass density profile of the membrane simulated with CHARMM36.

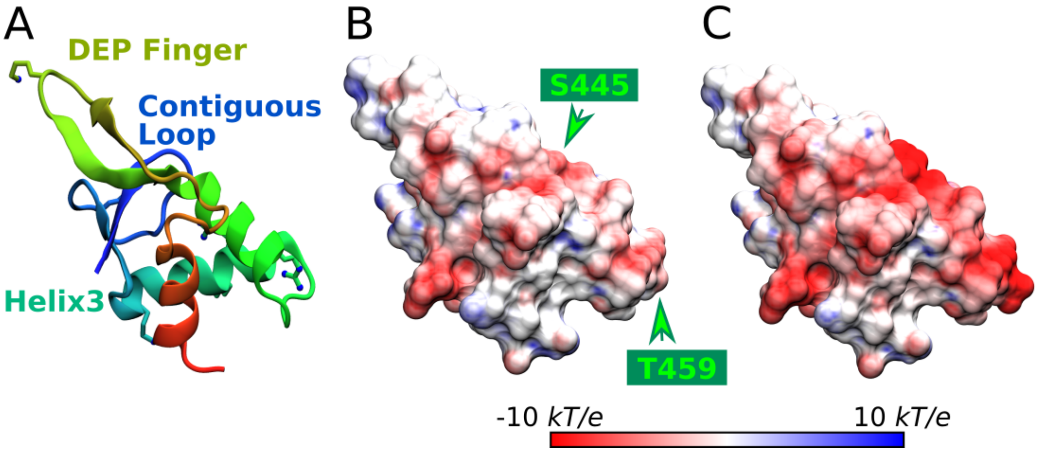


**Figure S3:** 180° rotation of Figure 1. (A): Cartoon representation of the domain highlighting the critical structural elements. The side chain of the positively charged residues involved in the membrane-binding are shown as balls and sticks. (B) and (C): Surface electrostatic potential for DEP-WT and DEP-P$^{[2-]}$. The arrows show the phosphorylation sites.

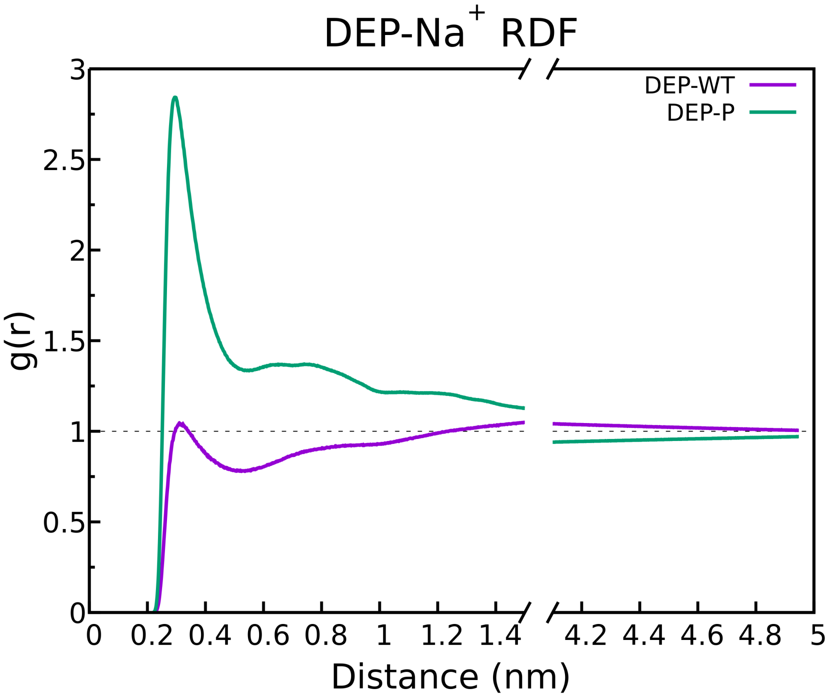


**Figure S4:** Radial Distribution Function (RDF) analysis of DEP domain and $Na^+$ from simulations using FAUNUS coarse-grained model. The peak for DEP-P indicates the strong interaction between the phosphorylated residues and the cations, resulting in effective screening of the negative charge.

**Table S6: Distribution of the per-residue distances from the membrane for the configurations populating the free energy minima in FAUNUS simulations of DEP-WT. Only the twenty closest residues are shown.**

| Residue[a] | Distance (Å) | |
|---|---|---|
| | Implicit Salt | Explicit Salt |
| W433 | 14.8 | |
| L434 | 14.0 | |
| K435 | 14.8 | |
| **R462** | | 13.1 |
| **R465** | 14.7 | 13.2 |
| **K466** | 13.9 | 11.8 |
| **S469** | 14.1 | 12.2 |
| **N470** | 14.9 | 12.7 |
| **L472** | 14.0 | 12.5 |
| **K473** | 13.0 | 11.2 |
| **A474** | | 13.6 |
| H479 | 13.2 | 12.5 |
| T480 | 14.9 | 14.6 |
| V481 | 11.9 | 11.6 |
| **N482** | 9.5 | 8.9 |
| **K483** | 10.0 | 9.1 |
| **I484** | 8.1 | 6.7 |
| **T485** | 9.4 | 7.9 |
| **F486** | 13.7 | 12.3 |
| **S487** | 13.0 | 11.9 |
| **E488** | | 14.5 |
| **Q489** | 13.8 | 13.1 |
| C490 | 14.7 | 14.1 |

[a] *Residues in bold belong to Helix 3 and the contiguous loop*

**Table S7: Cations Coordination Number of DEP Phosphosites**

| Simulations[a] | DEP Domain | pS445 | pT459 | pS469 | pT485 |
|---|---|---|---|---|---|
| Water + 150 mM NaCl | DEP-WT | 0.048 | 0.135 | 0.002 | 0.008 |
| | DEP-P[1-] | 1.102 | 1.061 | 0.030 | 0.104 |
| | DEP-P[2-] | 5.933 | 8.362 | 0.590 | 2.230 |
| POPA + 150 mM NaCl | DEP-WT | 0.710 | 3.796 | 0.129 | 0.291 |
| | DEP-P[1-] | 3.602 | 3.145 | 1.920 | 4.731 |
| | DEP-P[2-] | 5.465 | 9.923 | 4.876 | 8.154 |
| | DEP-WT (ECC) | 1.217 | 2.923 | 0.903 | 0.831 |
| | DEP-P[1-] (ECC) | 2.661 | 9.625 | 3.163 | 6.220 |
| POPA + 150 mM KCl | DEP-WT (ECC) | 0.349 | 0.602 | 0.203 | 0.514 |
| | DEP-P[1-] (ECC) | 0.863 | 0.862 | 0.669 | 1.744 |
| POPA + 10 mM $CaCl_2$ | DEP-WT (ECC) | 0.074 | 1.120 | 0.036 | 0.050 |
| | DEP-P[1-] (ECC) | 2.090 | 3.532 | 1.457 | 3.003 |
| POPA:POPC + 150 mM NaCl | DEP-WT | 0.216 | 1.505 | 0.013 | 0.141 |
| | DEP-P[1-] | 1.500 | 1.154 | 0.572 | 1.979 |
| | DEP-P[2-] | 9.510 | 7.891 | 0.584 | 6.865 |
| POPA[2-]:POPC + 150 mM NaCl | DEP-WT | 0.073 | 0.369 | 0.023 | 0.013 |
| | DEP-P[1-] | 0.625 | 5.529 | 0.929 | 2.333 |
| | DEP-P[2-] | 6.189 | 8.256 | 0.676 | 7.364 |

[a] *For DEP-membrane systems the values represent averages over both copies of the domain*

**Table S8: Hydrogen bonds of DEP domain and the phosphorylation sites proximal to the surface of POPA:POPC membrane, grouped by lipid type**

| | Hydrogen Bonds | | | | | |
|---|---|---|---|---|---|---|
| | DEP-WT | | DEP-P[1-] | | DEP-P[2-] | |
| | POPA | POPC | POPA | POPC | POPA | POPC |
| *Full domain* | 12.52 ± 3.16 | 3.58 ± 1.74 | 12.50 ± 4.43 | 3.17 ± 1.72 | 13.75 ± 3.44 | 2.51 ± 1.66 |
| *pT459* | 0.03 ± 0.17 | 0.00 ± 0.05 | 0.01 ± 0.07 | 0.00 ± 0.06 | 0.05 ± 0.22 | 0.00 ± 0.00 |
| *pS469* | 0.00 ± 0.05 | 0.00 ± 0.06 | 0.27 ± 0.53 | 0.00 ± 0.02 | 0.04 ± 0.19 | 0.00 ± 0.00 |
| *pT485* | 0.53 ± 0.62 | 0.14 ± 0.35 | 0.57 ± 0.67 | 0.13 ± 0.34 | 0.56 ± 0.60 | 0.00 ± 0.00 |

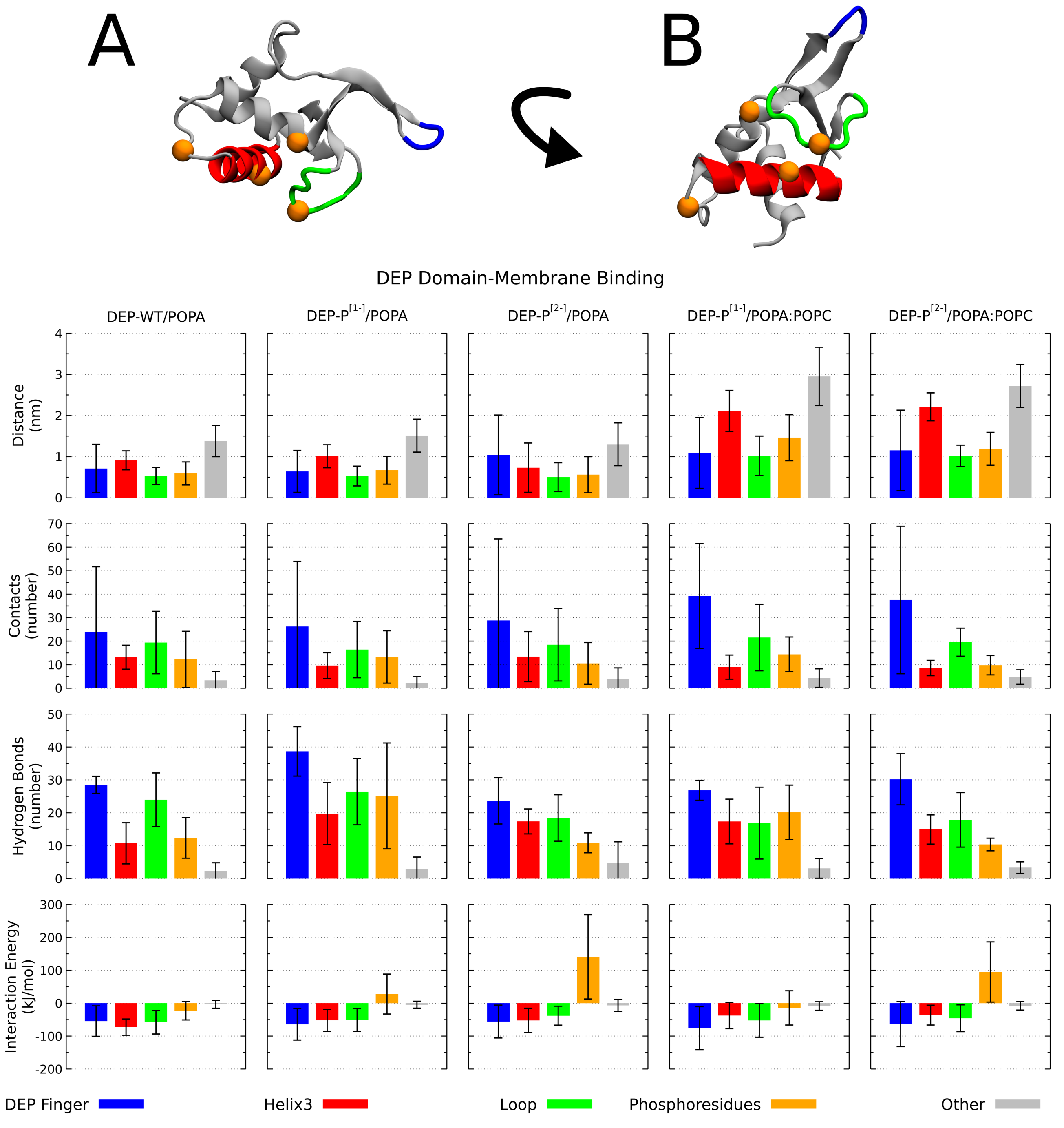


**Figure S5:** Protein regions critical for DEP-membrane binding. The location on the domain of the DEP finger (blue), Helix 3 (red), the contiguous loop (green), and the phosphoresidues (orange beads) is highlighted in panel A and B. For all mentioned regions, along with the remaining residues of DEP domain (in gray), the contacts, hydrogen bonds, interaction energy, and distance from the membrane (see Methods), are summarized in the bar plots. The large fluctuations associated with the DEP finger suggest a dispensable role in membrane binding, as result of its flexible nature. Similarly, not all phosphoresidues bind the membrane via ion-mediated interactions. On the contrary, Helix 3 and the contiguous loop are in general characterized by smaller fluctuations and strongly interacted with the membrane, regardless of the lipid composition. The shown systems were simulated with standard atomic charges and ~150 mM NaCl. The protonation state of the phosphate group on DEP domain is described by the superscripts. Note that all the values were normalized by the number of residues in each region and that for DEP-WT the corresponding unmodified amino acids are evaluated in place of the phosphoresidues.

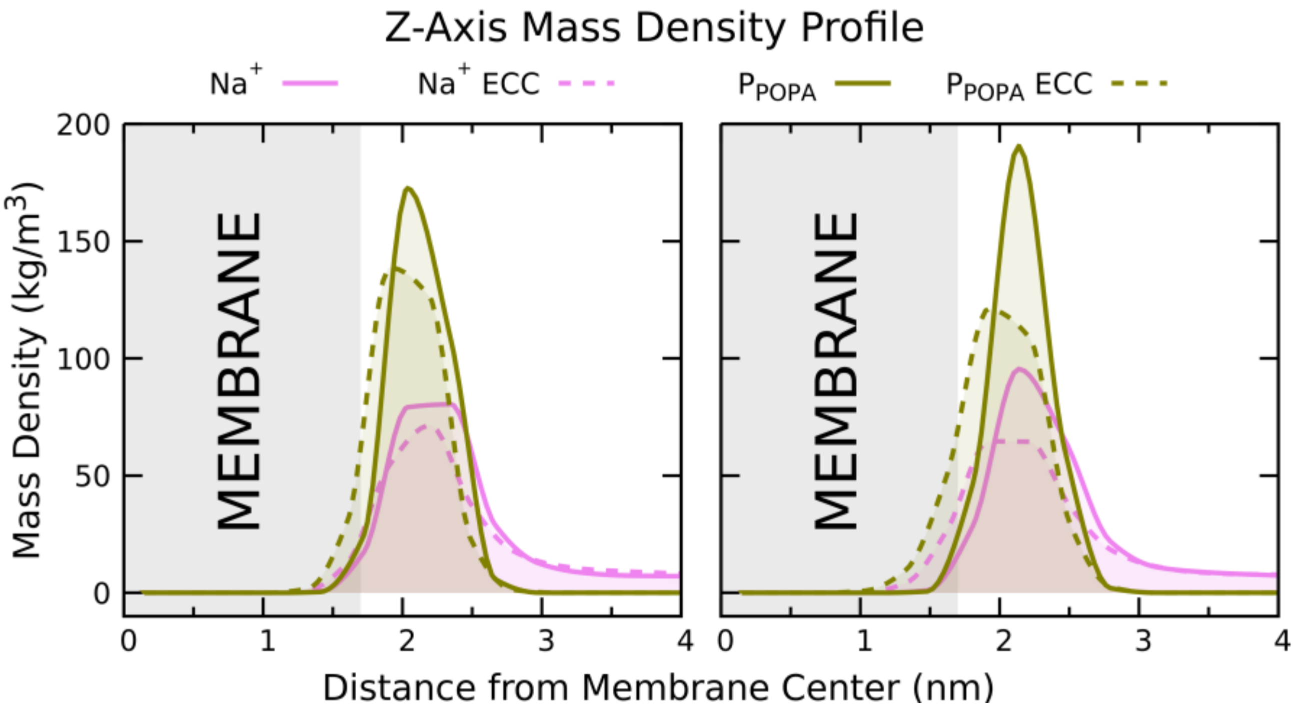


**Figure S6:** Averaged density profile along the membrane normal of $Na^+$ ions and P atom from POPA headgroup with standard and ECC charges for DEP-WT (left) and DEP-P$^{[1-]}$ (right). The profiles were symmetrized and show only half of the simulated box. The scaled charge scheme resulted in reduced accumulation of $Na^+$ ions at the membrane surface (pink dashed lines). Note that the ECC model caused looser headgroup packing and deeper penetration of the cations into the membrane (tan and pink dashed lines).

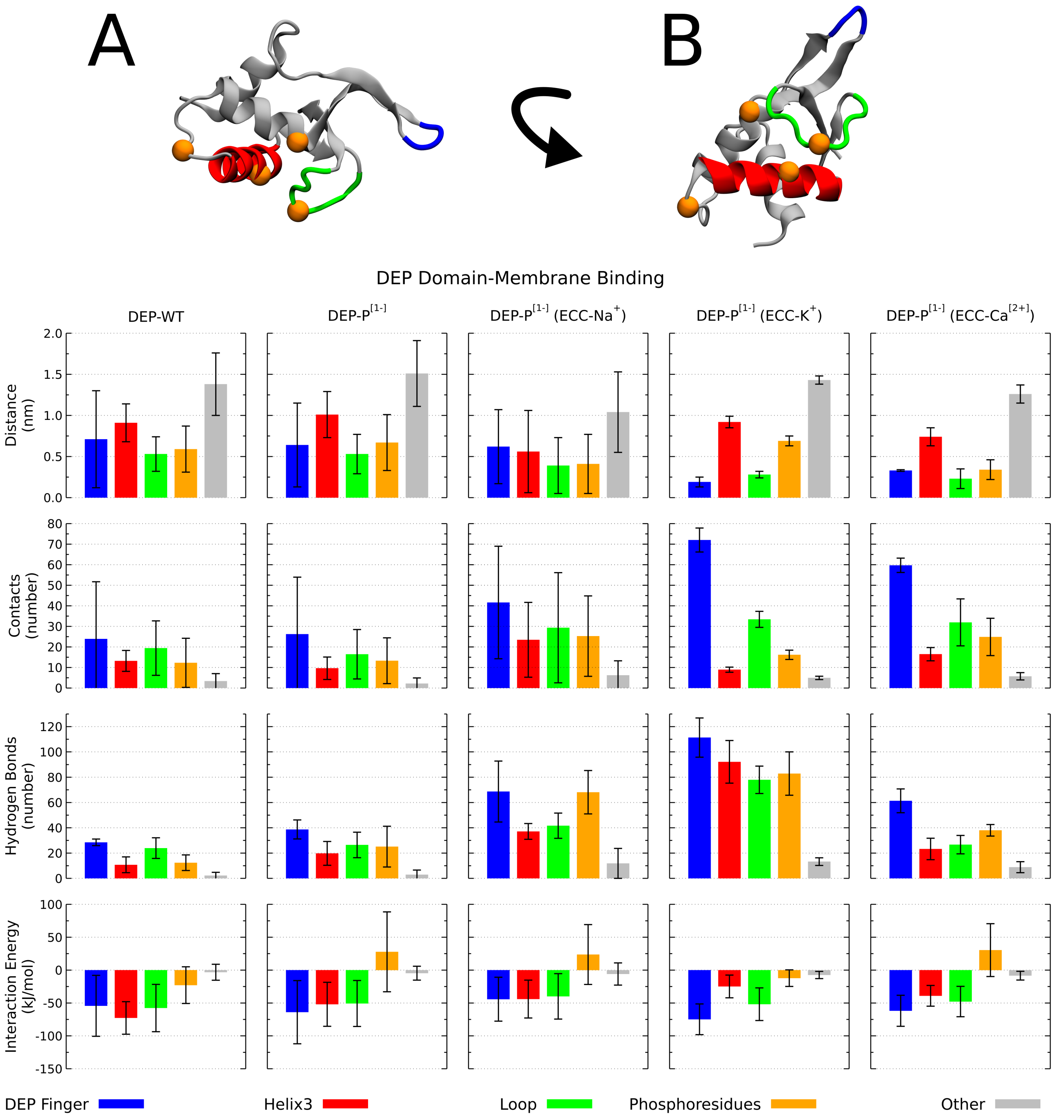


**Figure S7:** Protein regions critical for DEP-membrane binding. The location on the domain of the DEP finger (blue), Helix 3 (red), the contiguous loop (green), and the phosphoresidues (orange beads) is highlighted in panel A and B. For all mentioned regions, along with the remaining residues of DEP domain (in gray), the contacts, hydrogen bonds, interaction energy, and distance from the membrane (see Methods), are summarized in the bar plots. DEP-P[1-] bound the membrane through the same structural regions, irrespective of the used charge scheme and ion type. The smaller values of the distance from the membrane (first row) for the systems simulated with the ECC charges, suggest deeper penetration of the domain into the membrane. The shown systems were simulated at POPA membrane and ~150 mM NaCl/KCl or 10 mM $CaCl_2$. Note that all the values were normalized by the number of residues in each region and that for DEP-WT the corresponding unmodified amino acids are evaluated in place of the phosphoresidues.

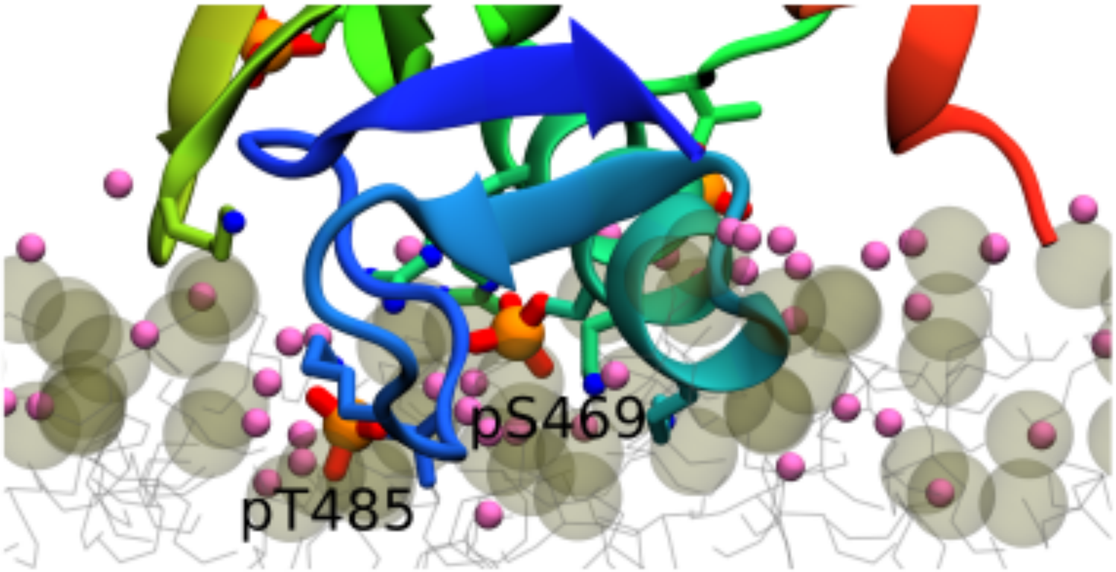


**Figure S8:** Final snapshot from 500 ns long simulation of DEP-P[1-] and ECC charge scheme. The ECC charges resulted in looser packing of the lipid molecules and deeper penetration of $Na^+$ ions and the phosphoresidues (e.g pT485 on the left) into the bilayer compared to simulations with standard atomic charges. Same color scheme and representation as in Figure 4.

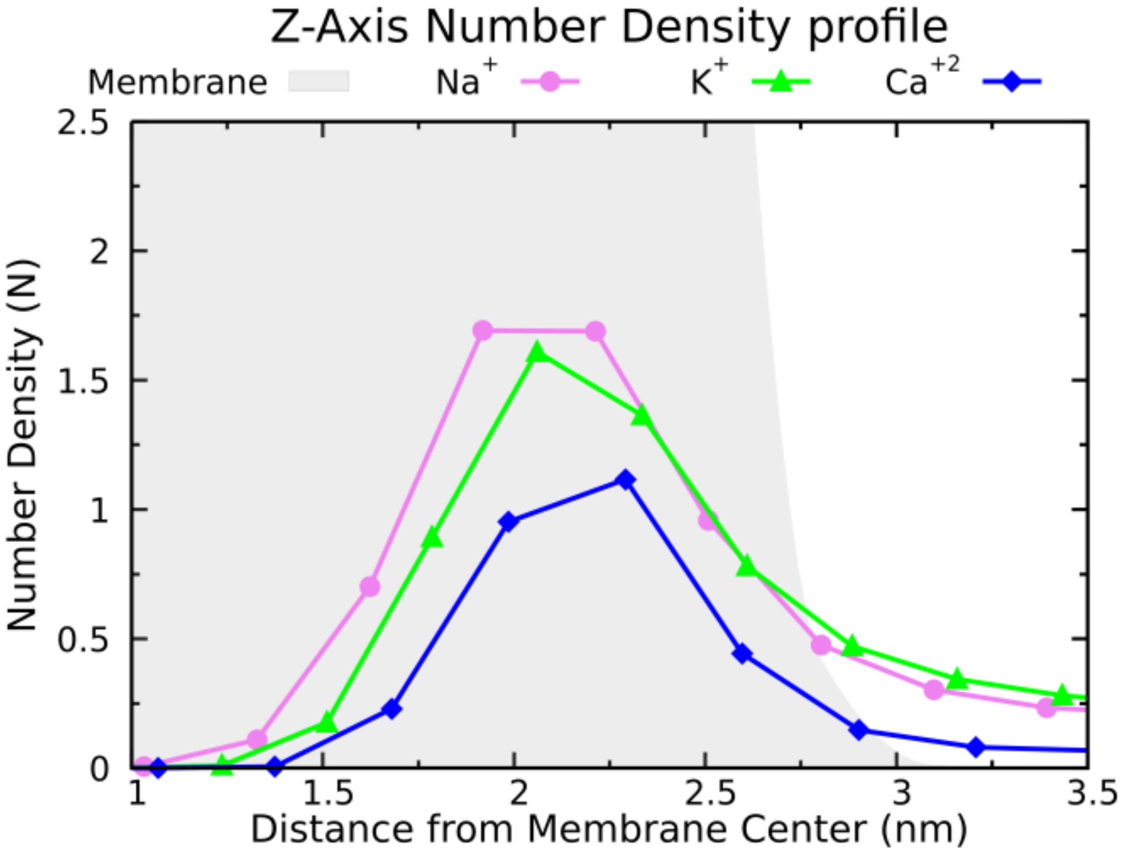


**Figure S9:** Averaged number density profiles along the membrane normal for $Na^+$, $K^+$ and $Ca^{2+}$ ions. The profiles were symmetrized and show half of the simulated box. Only the membrane density (gray area) from simulation with $Na^+$ is shown for simplicity. Note that the $Ca^{2+}$ concentration (10 mM) is lower than $Na^+$ or $K^+$ concentration (150 mM) by a factor of 15.

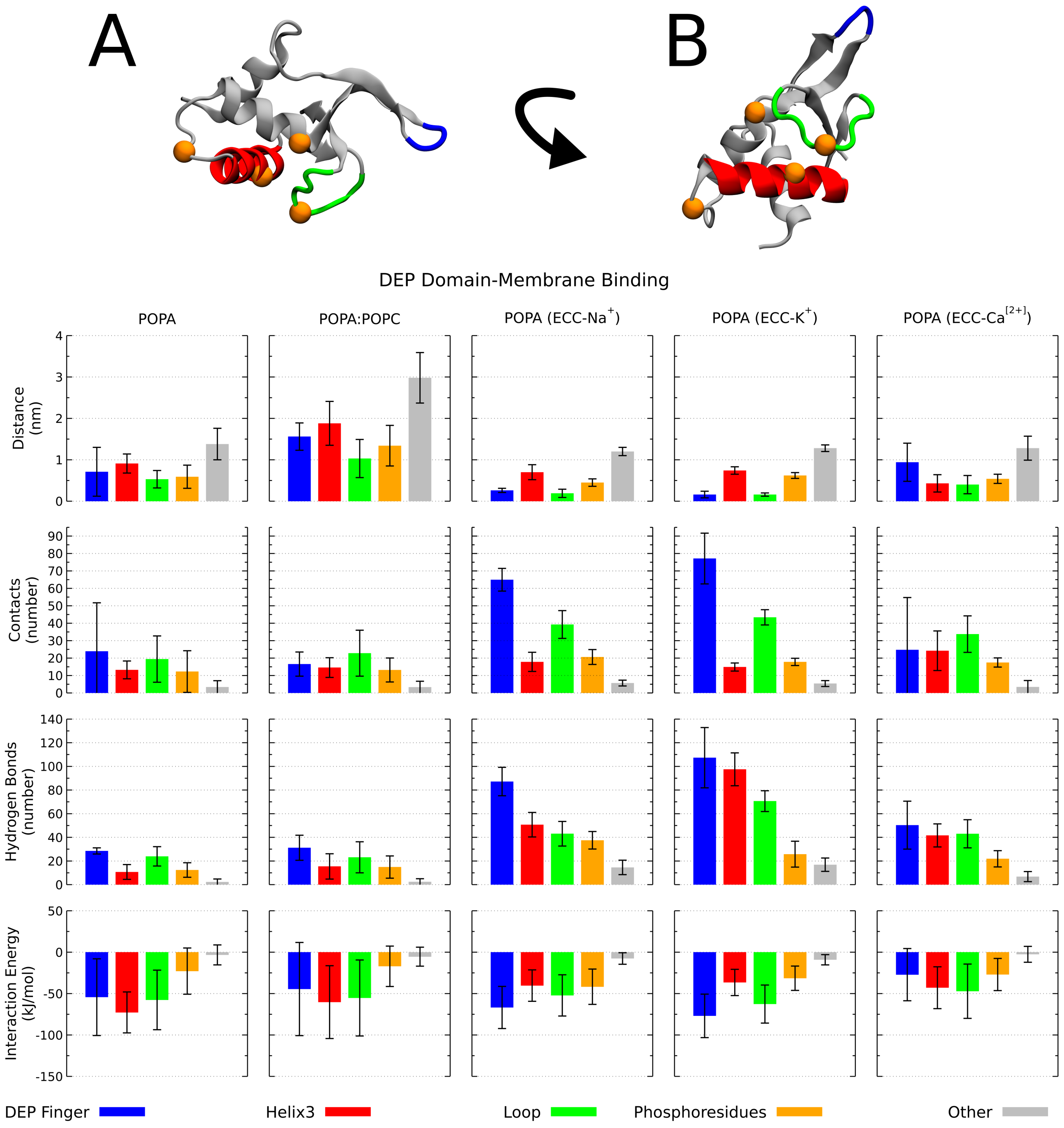


**Figure S10:** Protein regions critical for DEP-membrane binding. The location on the domain of the DEP finger (blue), Helix 3 (red), the contiguous loop (green), and the phosphoresidues (orange beads) is highlighted in panel A and B. For all mentioned regions, along with the remaining residues of DEP domain (in gray), the contacts, hydrogen bonds, interaction energy, and distance from the membrane (see Methods), are summarized in the bar plots. Irrespective of the used charge scheme, ion type, and lipid composition, the same structural regions of DEP-WT were involved in membrane binding. The smaller values of the distance from the membrane (first row) for the systems simulated with the ECC charged scheme, suggest deeper penetration of the domain into the membrane. The shown systems were simulated at ~150 mM NaCl/KCl or 10 mM $CaCl_2$. Note that all the values were normalized by the number of residues in each region and that the corresponding unmodified amino acids are evaluated in place of the phosphoresidues.

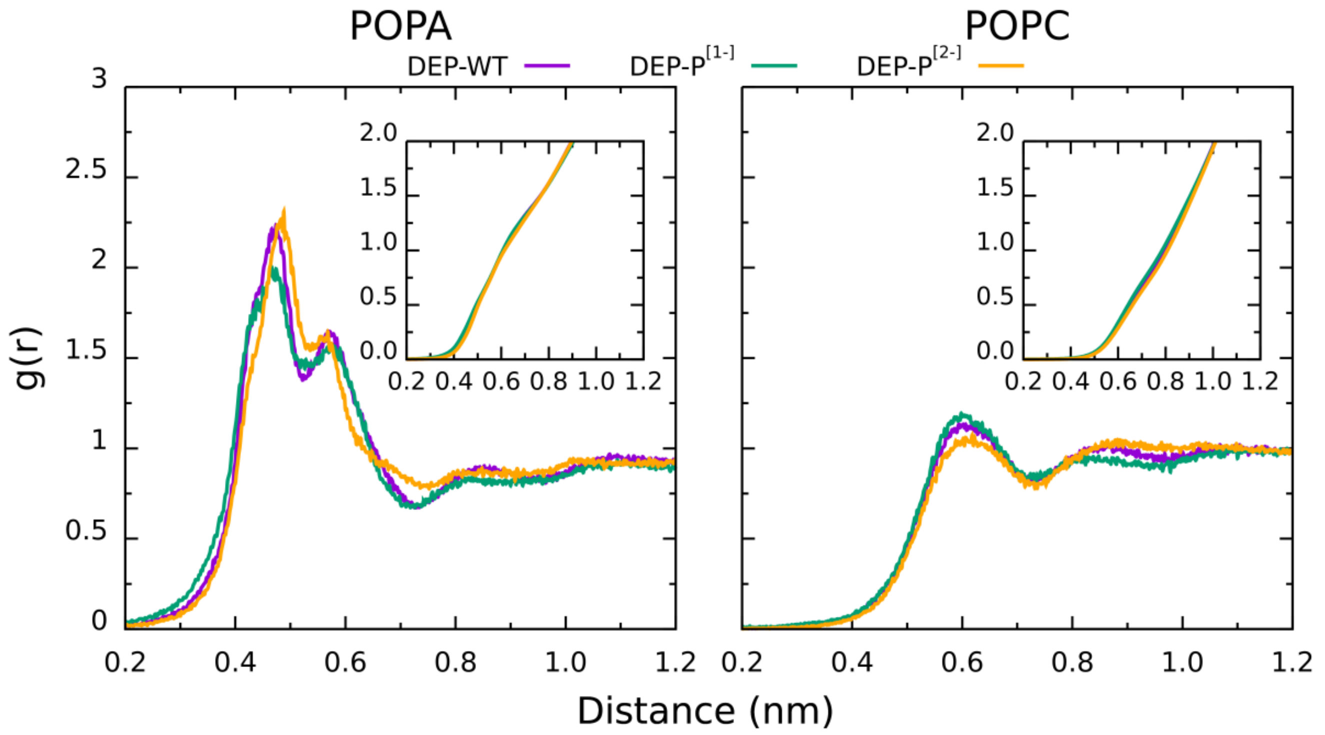


**Figure S11:** Radial distribution function (RDF) of all P atoms from POPA (left panel) and POPC (right panel) lipids for DEP-WT, DEP-P[1-], and DEP-P[2-] at POPA:POPC membrane. The cumulative number calculated as *int(g(r))* is displayed in the insets. The profiles were averaged over both lipid leaflets. No significant difference in the aggregation of both lipid species is observed in all simulated systems.

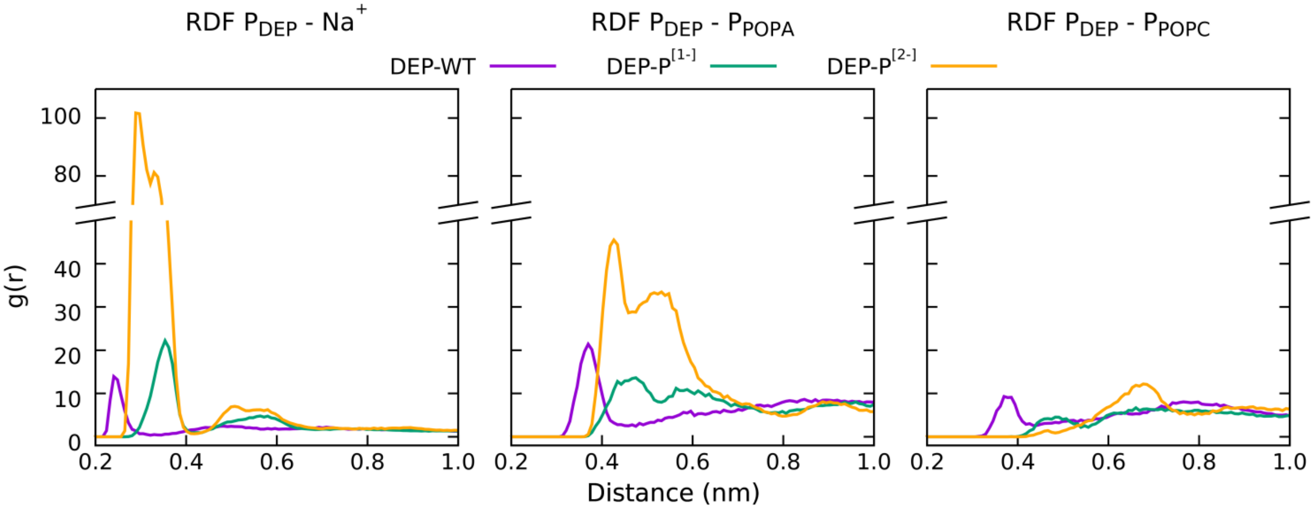


**Figure S12:** For DEP-WT, DEP-P[1-], and DEP-P[2-] at POPA:POPC membrane, the three panels show respectively the radial distribution function (RDF) between the P atoms from the domain and the Na[+] ions in solution (left), the P atoms from POPA (middle), and from POPC (right) lipids. The profiles were averaged over both lipid leaflets and copies of the domain. In the case of DEP-WT, the O atoms from the sidechain hydroxyl group were used. The doubly deprotonated phosphorylated residues of DEP are characterized by an higher ions coordination number, which could possibly lead to branched ion-mediated interactions. Note the similarities between the two-peaks profiles (orange lines) in the middle panel of this figure and the upper panel of Figure 6.

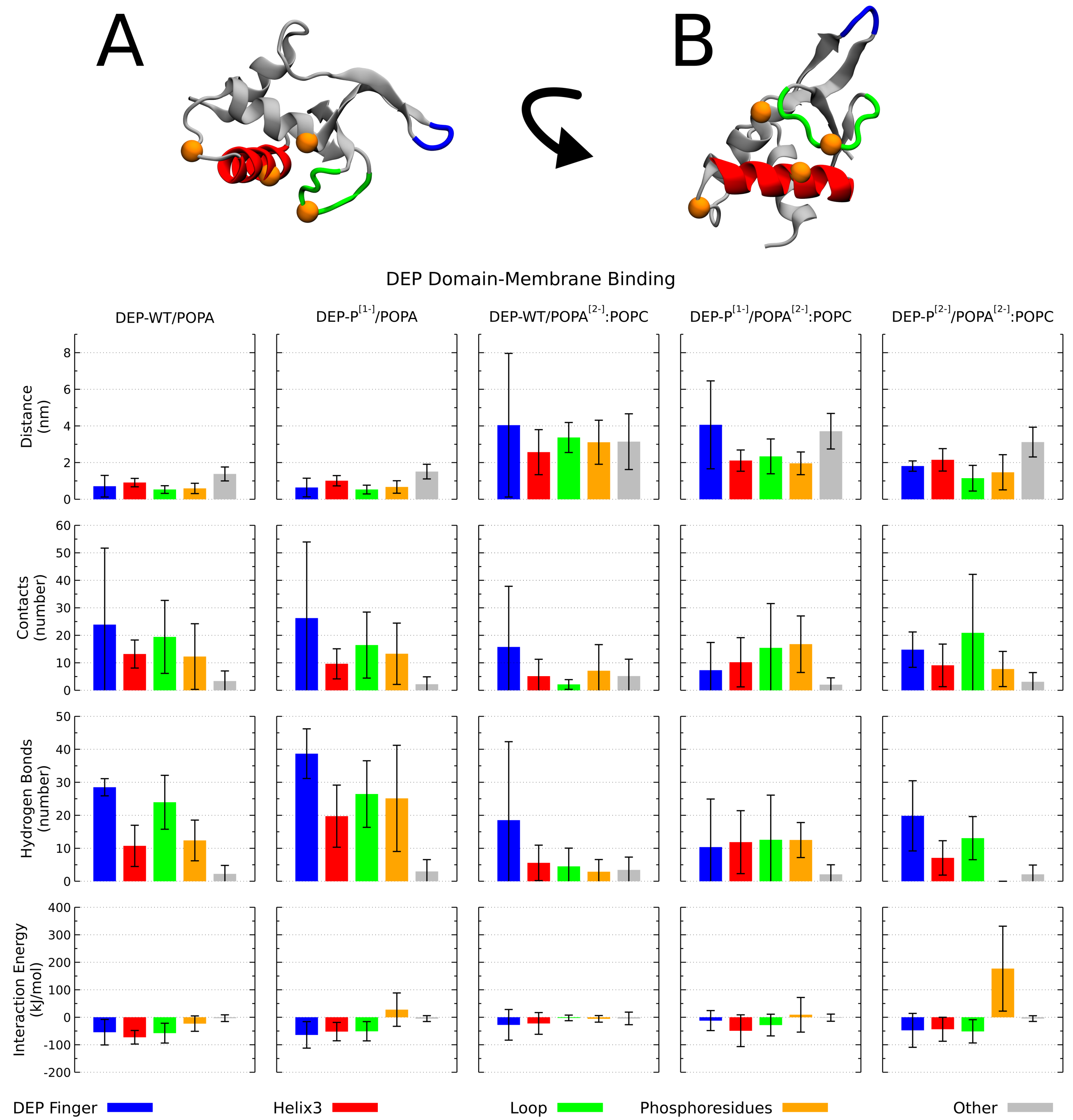


**Figure S13:** Protein regions critical for DEP-membrane binding. The location on the domain of the DEP finger (blue), Helix 3 (red), the contiguous loop (green), and the phosphoresidues (orange beads) is highlighted in panel A and B. For all mentioned regions, along with the remaining residues of DEP domain (in gray), the contacts, hydrogen bonds, interaction energy, and distance from the membrane (see Methods), are summarized in the bar plots. The strong charge density of doubly deprotonated PA led to strong binding of DEP domain. The different interaction pattern displayed by DEP-WT and DEP-P$^{[1-]}$ is indicative of a different binding orientation. Interestingly, DEP-P$^{[2-]}$ bound the membrane similar to DEP at full-PA membrane. The shown systems were simulated with standard atomic charges and ~150 mM NaCl. The protonation state of the phosphate group on DEP domain and PA lipids is described by the superscripts. Note that all the values were normalized by the number of residues in each region and that for DEP-WT the corresponding unmodified amino acids are evaluated in place of the phosphoresidues.

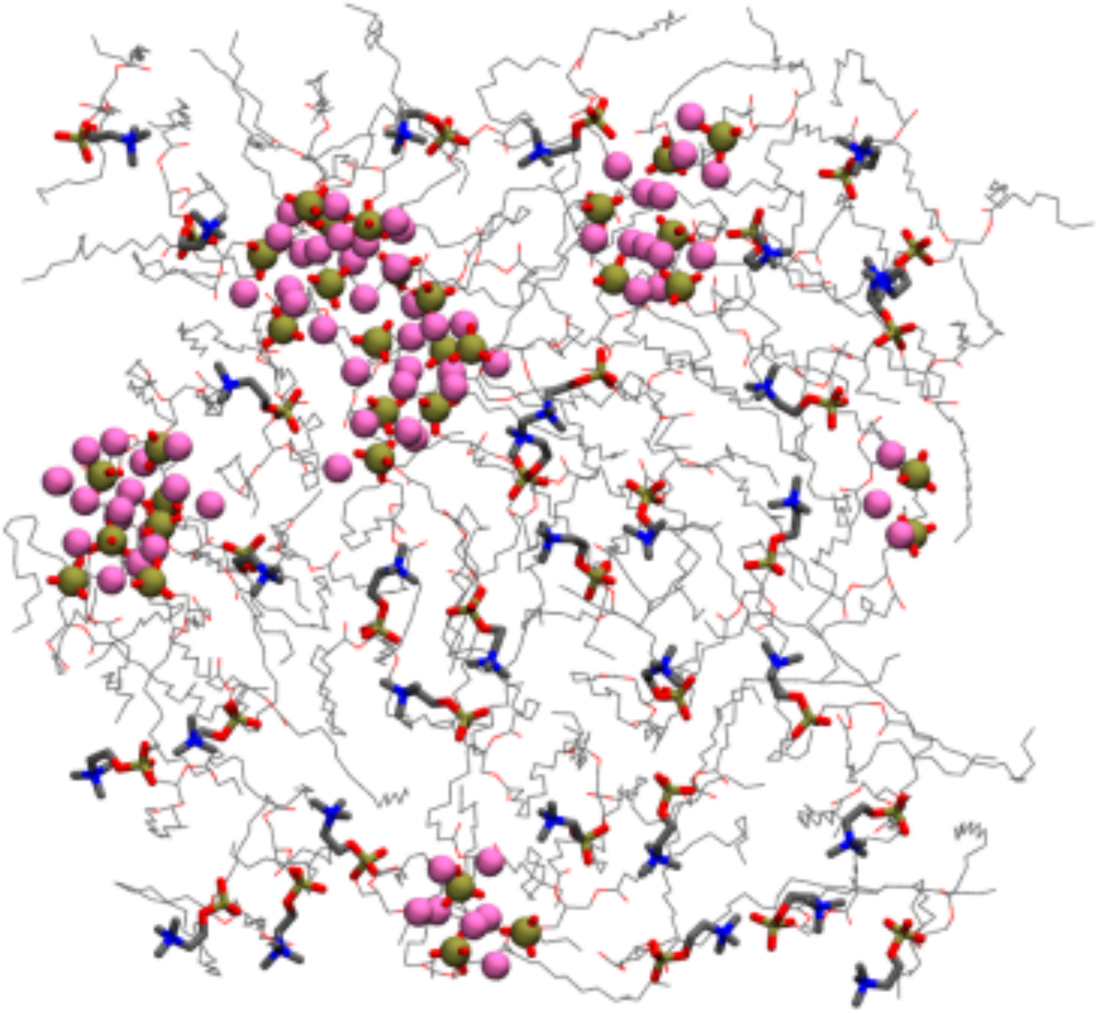

**Figure S14:** Top view of one leaflet from the final snapshot of 500 ns long simulation of DEP domain at membrane composed of POPC and doubly deprotonated POPA lipids at 1:1 ratio. POPA tends to form aggregates via cation-mediated interactions. For both lipid species, the glycerol and acyl tails are shown as lines, while the phosphates and headgroup substituents are displayed as spheres and cylinders. C, O, N, P, and $Na^{+}$ are colored in grey, red, blue, tan, and pink, respectively. P atoms from POPA and $Na^{+}$ ions are represented as VDW spheres, to highlight the aggregates. DEP domain, waters, and $Cl^{-}$ ions are omitted for clarity.

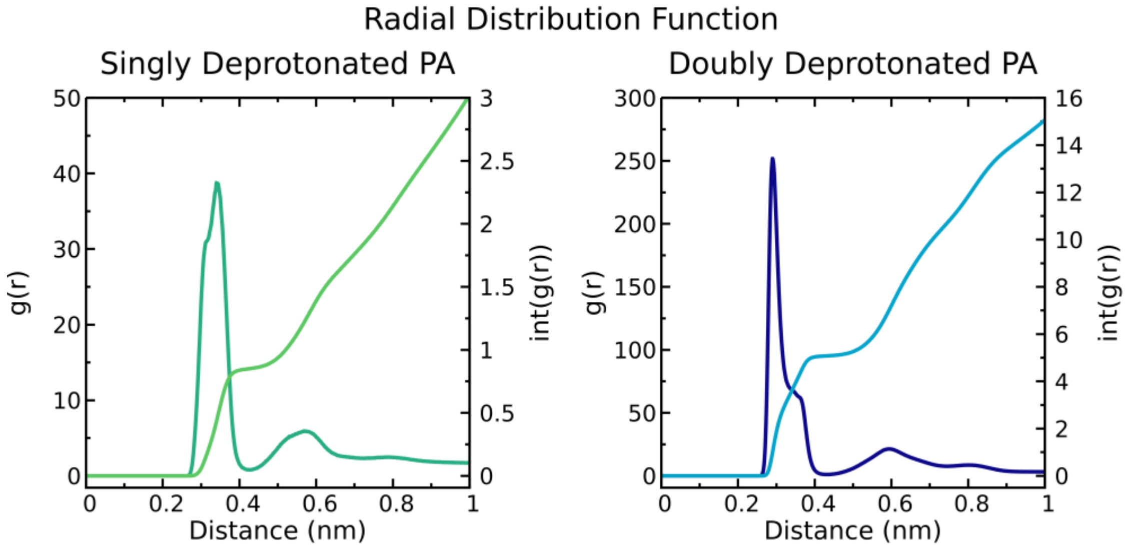


**Figure S15:** Radial Distribution Function (dark color) and cumulative number (light color) of P atom from singly (left panel) and doubly (right panel) deprotonated PA lipids and $Na^{+}$ ions.

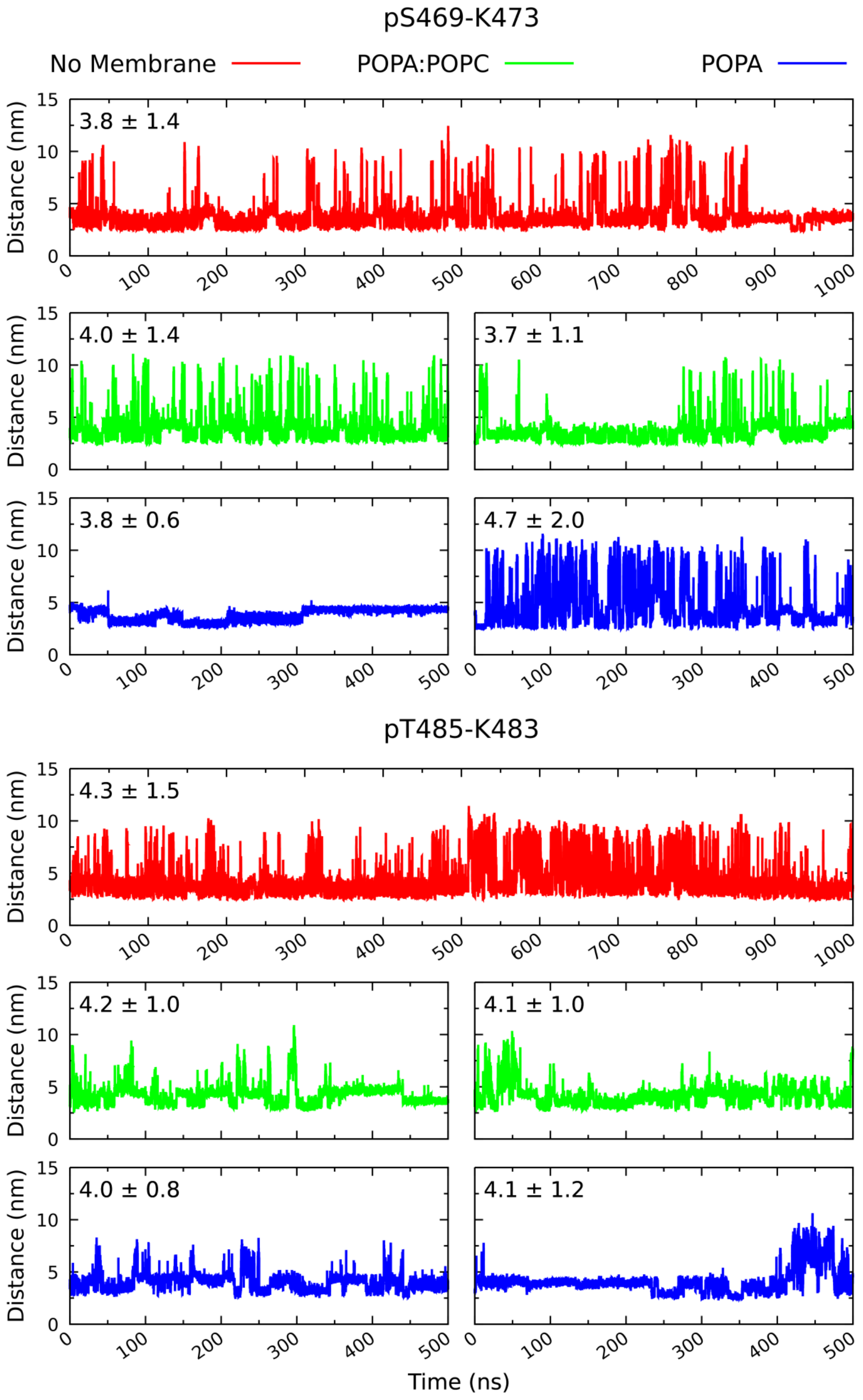


**Figure S16:** Time series of the salt bridges formed by doubly deprotonated pS469 (above) and pT485 (below) in simulations of DEP-P in solution (red), at POPA:POPC membrane (green), and at POPA membrane (blue). For the membrane bound domain both copies are shown. At the top-left of each plot is indicated the average and standard deviation of the length of the salt bridge. Note that, the intermolecular interaction involving pS469 is partially disrupted, while the one formed by pT485 is in general stabilized.

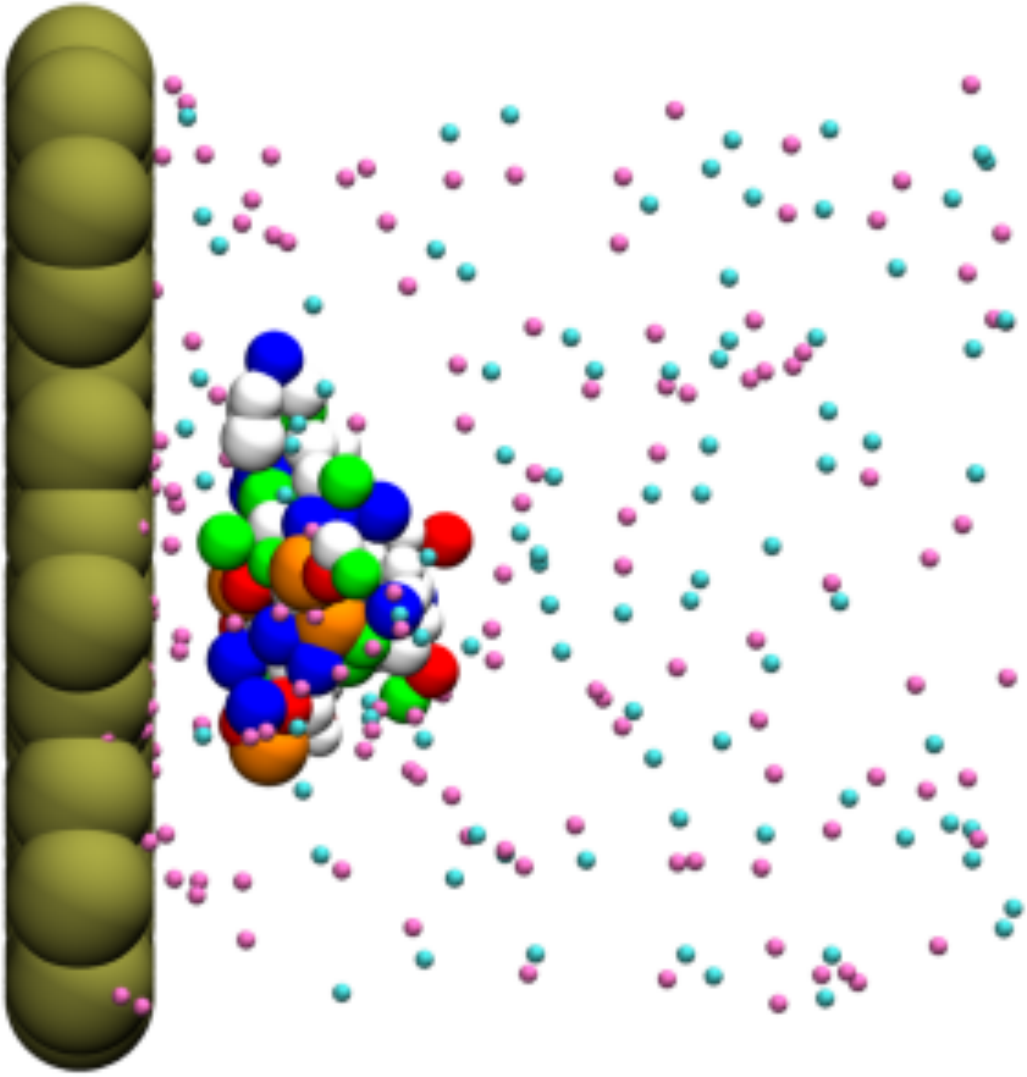

**Figure S17:** Alternative membrane binding mode of DEP-P from FAUNUS simulations with explicit ions. PA lipids are represented as tan beads, $Na^{+}$ are pink, and $Cl^{-}$ are cyan. Polar, apolar, basic, acidic, and phosphorylated residues are colored green, white, blue, red, and orange, respectively. Cations mediate the interactions between negatively charged phosphoresidues and PA lipids.
